# A LexA-like repressor and global H-NS-like regulators enable the fine-tuning of R-tailocin expression in environmental *Pseudomonas*

**DOI:** 10.1101/2025.01.21.634025

**Authors:** Clara Margot Heiman, Hammam Antar, Florian Fournes, Christoph Keel, Jordan Vacheron

**Author notes:** **Corresponding authors:** Christoph Keel and Jordan Vacheron, Department of Fundamental Microbiology, University of Lausanne, Biophore Building, CH-1015 Lausanne, Switzerland.

## Abstract

Bacteria rely on an arsenal of weapons to challenge their opponents in highly competitive environments. To specifically counter closely related bacteria, specialized weapons with a narrow activity spectrum are deployed, particularly contractile phage tail-like particles or R-tailocins. Their production leads to the lysis of the producing cells, indicating that their expression must be carefully orchestrated so that only a small percentage of cells produce R-tailocins for the benefit of the entire population. In this study, we set out to better understand how the production of these phage tail-like weapons is regulated in environmental pseudomonads using the competitive plant root colonizer and environmental model *Pseudomonas protegens* CHA0. Using an RNA sequencing (RNA-seq) approach, we found that genes involved in DNA repair, particularly the SOS response program, are upregulated following exposure of the pseudomonad to the DNA-damaging agents mitomycin C and hydrogen peroxide, while genes involved in cell division and primary metabolism are downregulated. The R-tailocin and prophage gene clusters were also upregulated in response to these DNA damaging agents. By combining reverse genetics, transcriptional reporters and chromatin immunoprecipitation sequencing (ChIP-seq), we show that the R-tailocin locus-specific LexA-like regulator PrtR1 represses R-tailocin gene expression by binding directly to the promoter region of the cluster, while the histone-like nucleoid structuring (H-NS) proteins MvaT and MvaV act as master regulators that indirectly regulate R-tailocin cluster expression. Our results suggest that at least these three regulators operate in concert to ensure tight control of R-tailocin expression and cell lytic release in environmental *Pseudomonas* strains.

**Author Summary:** To face their opponents in highly competitive environments, bacteria rely on an arsenal of weapons. To specifically target closely related bacterial strains, specialized weapons with a narrow activity spectrum are produced, notably contractile phage tail-like particles such as R-tailocins. Since production of these particles leads to the lysis and death of the producing cell, their expression must be tightly regulated. Here, we set out to better understand how the expression of these phage tail-like weapons is regulated in environmental bacteria using the competitive plant root colonizer and environmental model *Pseudomonas protegens* CHA0. We found that following DNA-damaging stress, which activates DNA repair systems and downregulates primary metabolism, the expression of gene clusters encoding viral-like particles such as the R-tailocins is upregulated. We identified cluster-specific and global regulators that are either directly or indirectly involved in the control of R-tailocin cluster expression. Our results contribute to a better understanding of the regulatory mechanisms of viral-like particle production in environmental *Pseudomonas* bacteria.

## Introduction

In highly competitive environments such as the plant rhizosphere or the animal gut, bacteria rely on an arsenal of weapons to counter their opponents. Although environmental bacteria produce broad-spectrum antimicrobials, these compounds are generally ineffective for competing against phylogenetically closely-related strains, as kin bacteria often share the same antimicrobials to which they are naturally immune (1–3). Alternatively, bacteria release compounds with a narrower activity spectrum to target kin strains. Among these weapons are contractile phage tail-like particles that include R-tailocins, which are highly specialized structures that specifically target only a small group of strains (1,4–10).

Like phages, these phage tail-like particles are encoded in dedicated prophage-like gene clusters in the genome of their host bacteria, but conversely to phages, these clusters no longer encode the capsid and the machinery necessary for replication (10). Similar to phages, the expression of R-tailocin gene clusters is induced upon activation of the bacterial SOS response to DNA damage (10). The SOS system has been thoroughly characterized in both *Escherichia coli* and *Bacillus subtilis*, where it alleviates the repression of more than 30 genes (11,12), while in *Pseudomonas aeruginosa* this system involves only 15 genes (13). Target genes in the SOS response are defined by their control by the two main regulators of this system, repression by LexA and activation by RecA (11,14). Under non-inducing conditions, homodimers of LexA, bound to cognate DNA sequences, the SOS boxes in the promoter region of the SOS response-related genes, repress the expression of target genes (14). Following the exposure to DNA-damaging agents such as mitomycin C (MMC) or ultraviolet rays (UVs), RecA binds to single-stranded DNA (ssDNA) to form activated RecA nucleoprotein filaments (RecA*) promoting DNA repair by homologous recombination (14). RecA* also acts as a coprotease to stimulate the serine protease activity of LexA, resulting in autocatalytic cleavage of LexA, leading to LexA depletion and the derepression of the SOS response genes (14,15). The SOS response includes activation of genes involved in DNA repair and cell division arrest, among others, as well as gene clusters encoding phages and phage tail-like particles (8,11–14,16–19).

The regulation of tailocin (pyocin) gene expression in response to DNA damage has been studied mainly in the human opportunistic pathogen *P. aeruginosa*. Two regulators, PrtN and the structurally LexA-related PrtR are involved (10,15,20). Under non-inducing conditions, the negative regulator PrtR represses the expression of *prtN*, the gene encoding a positive regulator of the tailocin gene cluster (20). Upon exposure to DNA-damaging agents, nucleoprotein filaments of RecA activate the autoproteolytic cleavage of PrtR, resulting in the de-repression of *prtN* (15). PrtN then binds to the P boxes present in the promoter region and activates the expression of the tailocin cluster genes (20). In laboratory practice, phages and phage tail-like particles are artificially induced with DNA-damaging agents known to trigger the SOS response, such as MMC, ciprofloxacin, reactive oxygen species (ROS), and UVs (10).

As these phage tail-like structures are used as weapons against bacterial competitors, global regulators involved in danger sensing or in virulence control could also be involved in their regulation and expression (21). As an example, the Gac/Rsm global regulatory pathway in *P. aeruginosa* has been shown to detect and respond to cell lysis induced by competitor bacteria by triggering the expression of competition factors such as the type VI secretion system (T6SS) or hydrogen cyanide for counterattack (22,23). Moreover, members of the histone-like nucleoid structuring (H-NS) protein family, are DNA-binding global repressors found in many Gram-negative bacteria that generally bind to AT-rich regions, allowing selective silencing of xenogeneic DNA, i.e. DNA acquired from foreign sources such as through horizontal gene transfer (24–28). It has been suggested that the cell then evolves control systems such as the recruitment of new positive regulators to take advantage of the newly acquired genes (27). *Pseudomonas* strains possess two such xenogeneic silencers, termed MvaT and MvaU in *P. aeruginosa* (29), or MvaT and MvaV in *Pseudomonas protegens* (30). In *P. aeruginosa*, these H-NS-like proteins function coordinately to control the expression of genes involved in virulence, cell surface structuring, housekeeping functions, and quorum sensing (29,31–36). These proteins are also involved in the silencing of prophage and pyocin S gene expression in *P. aeruginosa* (33,37,38). In *P. protegens*, they have been shown to globally affect the production of the antimicrobials pyoluteorin and 2,4-diacetylphloroglucinol (DAPG), exoenzyme activity, cell surface properties, motility, and biocontrol capabilities against a plant pathogen (30). However, their role in tailocin regulation has not been investigated yet.

The root-colonizing bacterium *P. protegens* type strain CHA0 (hereafter referred to as CHA0) used in this study is a widely used environmental model strain (1,39,40). The genome of CHA0 harbors a gene cluster encoding two R-tailocins along with the cell lytic enzymes required for their explosive release into the extracellular environment (1). Additionally, the genome contains two prophages: an active siphovirus and a myovirus that is thought to be cryptic (1,10). In contrast to *P. aeruginosa*, the R-tailocin gene cluster of CHA0, as well as those of other environmental *Pseudomonas* species, encodes only the negative regulator PrtR1, while PrtN is absent (1,41,42). Since the regulation of R-tailocin expression in environmental pseudomonads has not been studied in detail, we aimed to identify the regulatory mechanisms governing R-tailocin gene cluster expression in this model strain. First, we performed RNA sequencing (RNA-seq) on CHA0 cultures exposed or not to MMC or hydrogen peroxide (H2O2). As expected, we found that under inducing conditions, genes involved in cell division and primary metabolism were downregulated, while those involved in DNA repair, particularly the SOS system, were upregulated. Accordingly, the R-tailocin and prophage clusters were also upregulated in response to these DNA damaging agents. We then investigated the R-tailocin locus-specific LexA-related regulator PrtR (termed PrtR1 due to the presence of two additional LexA-related PrtR proteins encoded on the CHA0 chromosome) and the two H-NS-like global regulators MvaT and MvaV, using transcriptional reporters in different mutant backgrounds, chromatin immunoprecipitation sequencing (ChIP-seq) and RNA-seq of a mutant lacking both *mvaT* and *mvaV*. We found that PrtR1 binds specifically to the promoter region of the R-tailocin gene cluster of CHA0. In contrast, MvaT and MvaV do not bind to the promoter regions of either *prtR1* or the R-tailocin gene cluster but instead bind broadly to various regions across the CHA0 chromosome. Nevertheless, mutants lacking both *mvaT* and *mvaV* displayed different R-tailocin gene expression patterns compared to the wild type, suggesting that the H-NS like regulators influence R-tailocin gene cluster expression indirectly.

## Results

### DNA-damaging compounds dysregulate the primary metabolism and induce DNA damage repair systems prompting viral particle expression

An RNA-seq approach was used to identify genes of *P. protegens* CHA0 that are influenced upon exposure of the strain to the DNA-damaging agents MMC and H2O2. The two agents are known to trigger the bacterial SOS system and the production of viral particles, including tailocins (8,10,15). They were chosen as MMC is a potent inducer of the SOS response and is widely used to induce the production of viral particles, while H2O2 is more representative of stresses found in natural environments. We used 9 µM of MMC, a concentration used in previous work (1), which allows the monitoring of R-tailocin gene expression in CHA0 without directly killing the cells before they produce viral particles. We identified 10 mM of H2O2 as a suitable concentration for tailocin monitoring, as it delayed bacterial growth and strongly induced R-tailocin gene expression (measured with the transcriptional reporter pOT1e-P*hol*-*egfp* in CHA0, **S1 Table**), similar to the effect of 9 µM MMC (**S1 Fig**). At lower levels of H2O2, R-tailocin gene expression was minimal or delayed, while at the highest concentration tested, cell growth was inhibited (**S1 Fig**).

For RNA-seq analysis, RNA was harvested from CHA0 cells at 3 h post induction with either 9 µM MMC or 10 mM H2O2 and sequenced (**S1 Fig**). We observed differential expression of genes involved in the formation and release of viral particles in CHA0, depending on the DNA-damaging agent used. Genes related to the production of the two R-tailocins and the *Siphoviridae* prophage in the CHA0 genome (1) were highly induced upon MMC exposure (**Fig 1A**). While the R-tailocin genes were also highly expressed in the presence of H2O2, the siphovirus genes showed lower expression (**Fig 1B**). Genes associated with the *Myoviridae* prophage, the second intact prophage in the CHA0 genome, were not induced by either H2O2 or MMC (**Fig 1A**, **1B**), supporting the hypothesis that this prophage is cryptic (1). Interestingly, the expression of the *prtR1* gene, which encodes the predicted negative regulator of the R-tailocin gene cluster, was upregulated in the MMC condition, but not in the H2O2 condition (**Fig 1A**, **1B**). Additionally, PPRCHA0_2151, a gene located in a defective prophage region and encoding a lectin-like bacteriocin homologous to LlpA2 of the closely related *P. protegens* strain Pf-5 (43), was also strongly upregulated in both conditions (**Fig 1A**, **1B**).

**Figure 1.**
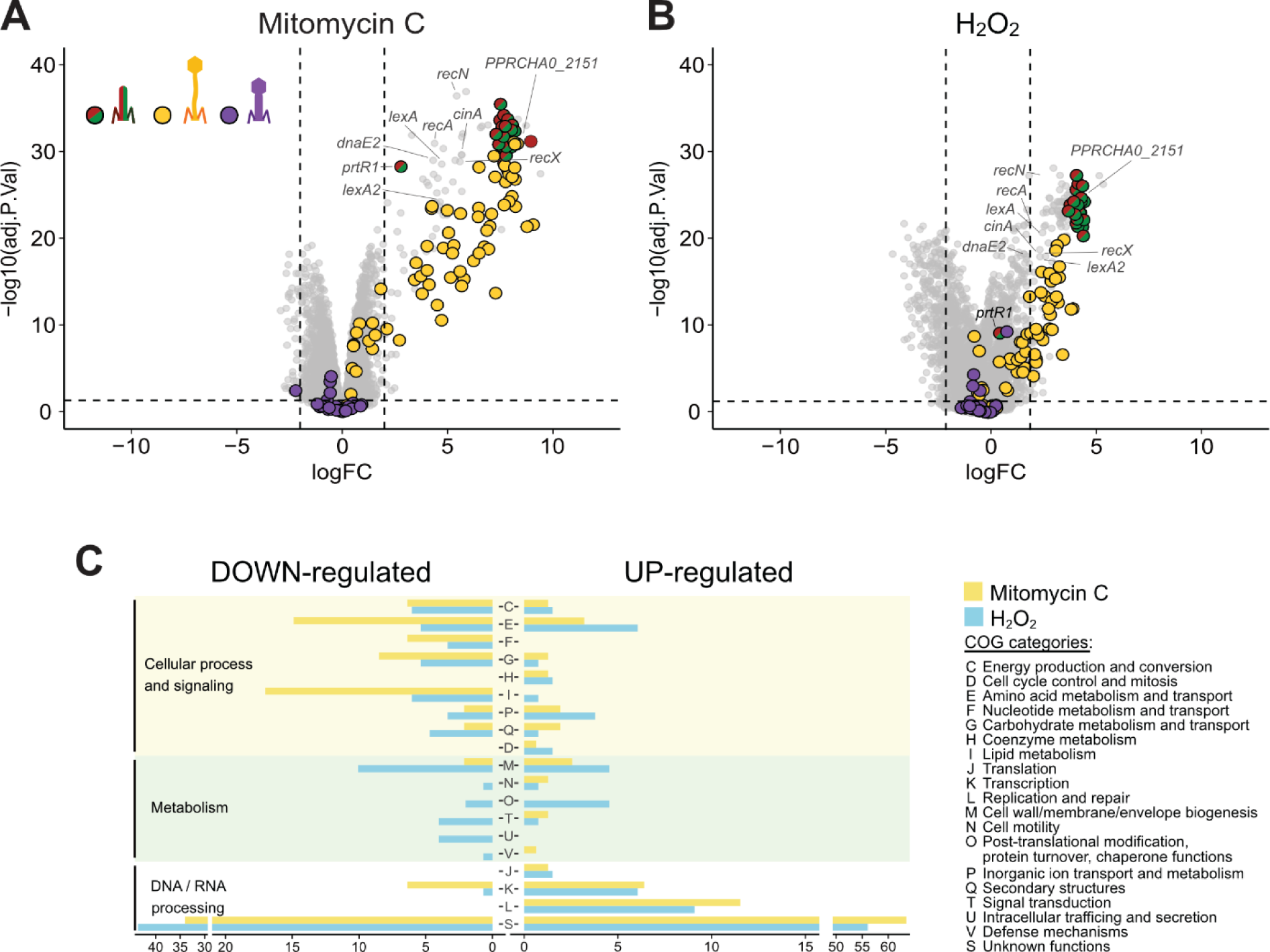
Mitomycin C and H2O2 differentially induce and repress the transcriptome of *P. protegens* CHA0. (**A**, **B**) Volcano plots showing the RNA sequencing results of CHA0 following exposure to 9 µM mitomycin C (**A**) or 10 mM H2O2 (**B**). The colored dots correspond to viral particle-associated genes, red and green for the two tailocins, yellow for the siphovirus, and purple for the myovirus of *P. protegens* CHA0. The vertical dashed lines correspond to the log2(Fold-change) thresholds set for this analysis (-2 >log2(FC) > 2). The horizontal dashed line corresponds to the significance level of *P* < 0.05. (**C**) Down-regulated and up-regulated genes according to their COG assignments following exposure to the two compounds. The length of the bars corresponds to the percentage of genes associated with the different COG assignments.

Differences for genes involved in the bacterial SOS response were detected when comparing the MMC and H2O2 conditions with the non-induced control. In the presence of both MMC and H2O2, genes such as *recN*, *cinA*, *recA*, *recX*, *dnaE2* (PPRCHA0_3719, DNA polymerase III, alpha subunit) as well as both *lexA* and *lexA2,* were upregulated (**Fig 1A**, **1B**).

Differentially regulated genes in the two conditions (MMC and H2O2) compared to the non-induced control were categorized into Clusters of Orthologous Genes (COG) assignments (**Fig 1C**). In response to the DNA-damaging agents, CHA0 cells downregulated genes associated with primary metabolism, including those involved in acid metabolism and transport, carbohydrate metabolism and transport, lipid metabolism, and cell wall / membrane envelope biogenesis (COG assignments E, G, I, and M respectively, **Fig 1C**). Conversely, genes related to transcription, replication and repair (COG assignments K and L, respectively) were upregulated (**Fig 1C**). There were differences in gene categorization depending on the agent. In the presence of H2O2, CHA0 cells downregulated more genes involved in cell wall / membrane envelope biogenesis (COG assignment M) and upregulated more genes related to amino acid metabolism and transport and transcription (COG assignments E and K, respectively) compared to MMC-exposed cells (**Fig 1C**). In contrast, MMC-exposed CHA0 cells had more downregulation of genes associated with cell wall / membrane envelope biogenesis (COG assignment M) and more upregulation of genes related to transcription and replication and repair (COG assignments K and L, respectively) compared to cells exposed to H2O2 (**Fig 1C**). In addition, the growth of CHA0 was less impacted by H2O2 than by MMC (**S1 Fig**).

Together, these results demonstrate that different DNA-damaging compounds can induce genes associated with the SOS response, and viral particle formation and release, in CHA0. Depending on the nature of the compound, the DNA-damaging agents affect CHA0 differently, resulting in different stress responses and differences in gene regulation and expression.

### Control of R-tailocin gene expression in *P. protegens* involves direct inhibition by the locus-specific regulator PrtR1

To better understand the regulation of the R-tailocin gene cluster of CHA0, we first aimed at understanding how the LexA-like locus-specific regulator PrtR1 interferes with R-tailocin expression, as PrtR proteins have been shown to control the expression of tailocins (pyocins) in *P. aeruginosa* (8,10,20). To do so, we monitored the expression of transcriptional reporters for the R-tailocin gene cluster (pOT1e-P*hol*-*egfp*) and the *prtR1* gene encoding the predicted negative regulator of the R-tailocin gene cluster (pOT1e-P*prtR1*-*egfp*) in different CHA0 mutant backgrounds under MMC-induced and non-induced conditions (**Fig 2A**, **S2 Fig**). Importantly, in our assays, the deletion of the PrtR1 regulator itself was not viable. Indeed, we expected PrtR1 of CHA0 to be a negative regulator of R-tailocin expression, analogous to PrtR previously described for *P. aeruginosa* (20). The removal of this gene would result in the constitutive expression of the R-tailocin structural genes as well as the genes encoding lytic enzymes to release the R-tailocins leading to fatal cell lysis. To circumvent this problem, we first deleted the entire R-tailocin gene cluster including all the lytic enzymes (holin and endolysins) (Δtailcluster) and then removed the *prtR1* gene, leading to the construction of a “conditional” mutant of *prtR1* (Δ*prtR1\**; **S2 Table**). As a control, we also tested our reporters in the Δtailcluster background. The CHA0 derivatives were exposed or not to 9 µM MMC, and the OD600nm as well as the GFP fluorescence were monitored for 24 h to assess the effect of the DNA-damaging compound on the expression of the different reporter constructs (**Fig 2A**, **S2 Fig**).

**Figure 2.**
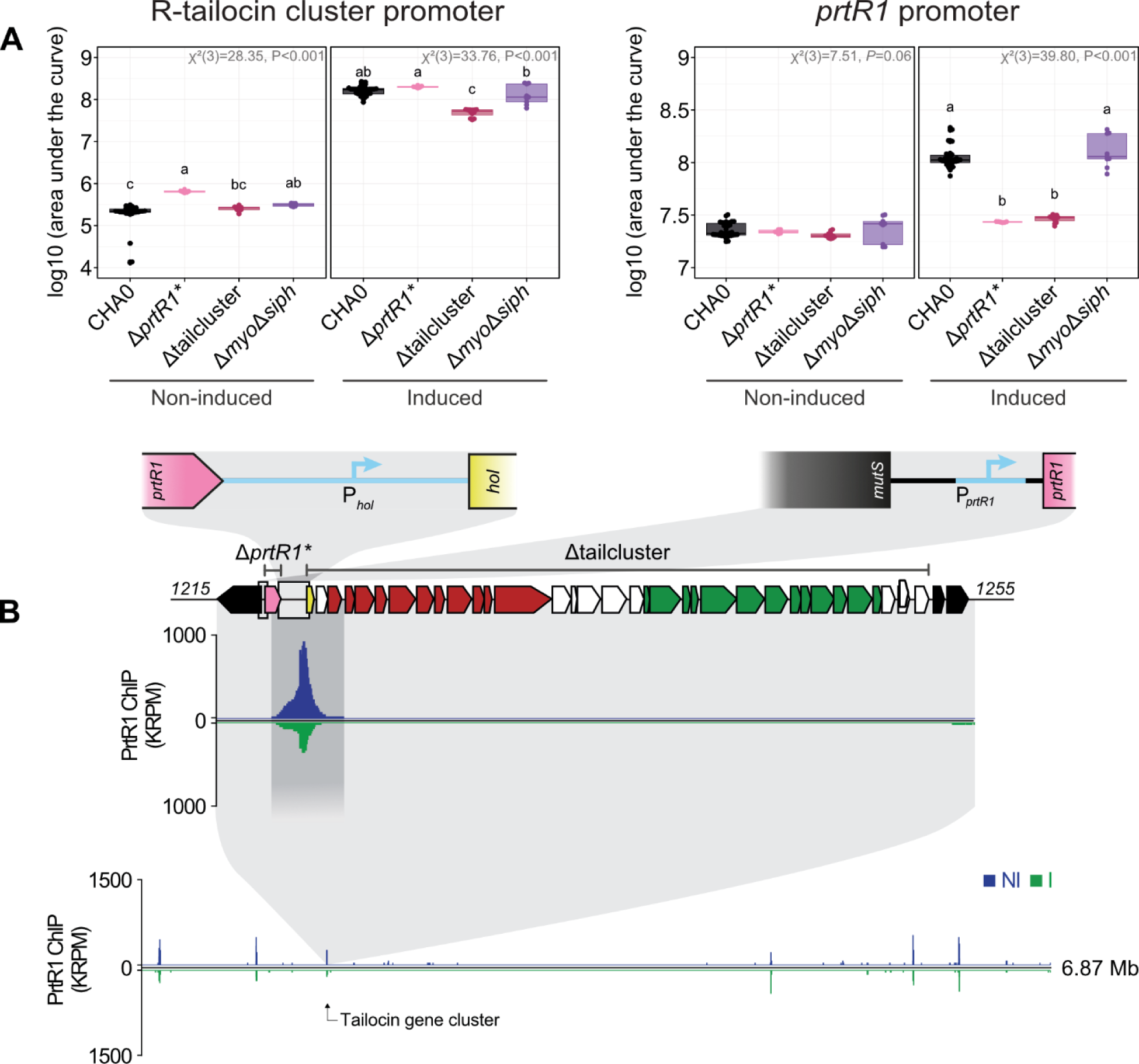
PrtR1 directly inhibits the expression of the R-tailocin gene cluster in CHA0 by interacting with the upstream region of the cluster. (**A**) The expression of the R-tailocin gene cluster and the locus-specific regulatory gene *prtR1* was monitored using transcriptional reporters pOT1e-*Phol-egfp* and pOT1e-P*prtR1-egfp*, respectively, in the wild type CHA0 (black) and its mutants Δ*prtR1\** (pink) and Δtailcluster (burgundy). Strains were grown in rich medium (NYB) following induction with 9 µM mitomycin C or without induction. Optical density at 600 nm and GFP fluorescence (relative fluorescence units, RFU) were monitored every 10 min for 24 h in a BioTeK Synergy H1 plate reader. Detailed curves are shown in **S2 Figure**. The data were then used to calculate the area under the RFU curve. Statistical differences were assessed by Kruskal-Wallis tests with Bonferroni correction and are indicated by letters. A minimum of three biological replicates with at least three technical replicates are plotted. (**B**) ChIP-seq results over the R-tailocin gene cluster of CHA0 for PrtR1 under non-induced (NI, blue) and induced (I, 9 µM mitomycin C, green) conditions. The genes encoding the structural parts of the R-tailocin #1 and the R-tailocin #2 are colored red and green, respectively, the *prtR1* gene is colored pink, the gene encoding the lytic gene holin (*hol*) is colored yellow, other genes belonging to the cluster are colored white, and the bacterial genes neighboring the R-tailocin gene cluster are colored black.

We observed that the expression of the R-tailocin gene cluster was approximately 3-log fold enhanced in the MMC-induced condition compared to the non-induced condition in the wild type CHA0 as well as the two mutant backgrounds Δ*prtR1\** and Δtailcluster (**Fig 2A**). However, the basal R-tailocin gene cluster expression was higher in the Δ*prtR1\** mutant compared to the wild type CHA0 (**Fig 2A**). This is specific to the *prtR1* deletion as the Δtailcluster mutant had similar expression levels as the wild type, supporting that PrtR1 acts as a negative regulator of R-tailocin expression in CHA0 but also suggesting the involvement of other important mechanisms of regulation (**Fig 2A**).

Furthermore, we investigated if PrtR1 influences its own expression. The expression of *prtR1* in the non-induced condition was the same in all three genetic backgrounds (**Fig 2A**). However, *prtR1* expression was significantly decreased in the MMC-induced condition in the two mutant backgrounds (Δ*prtR1\** and Δtailcluster) compared to the wild type CHA0, suggesting that a potential positive regulator of *prtR1* is encoded within the R-tailocin gene cluster (**Fig 2A**). We therefore tested the expression of *prtR1* in CHA0 mutants deleted for suspected regulators within the R-tailocin gene cluster, i.e. Δ*lateD*, Δ*PPRCHA0_1250*, Δ*PPRCHA0_1251* and Δ*PPRCHA0_1252*, but did not detect any marked differences in either non-induced or induced conditions (**S3 Fig**). Moreover, to identify if there was a possible crosstalk between the other viral particle-related gene clusters present in the CHA0 genome, we measured the expression of the R-tailocin gene cluster and of *prtR1* in a mutant lacking both the *Siphoviridae* and *Myoviridae* prophages (Δ*myo*Δ*siph*). We showed that the removal of these two clusters did not impact the expression of either the R-tailocin cluster or the *prtR1* gene in both non-inducing and inducing conditions (**Fig 2A, S2 Fig**), suggesting a high specificity of viral-like regulators.

To confirm that PrtR1 binds to the R-tailocin gene cluster in CHA0, we performed a chromatin immunoprecipitation sequencing (ChIP-seq) analysis. We used a mutant that produces the PrtR1 protein with a C-terminal V5-tag expressed from the endogenous locus (**S2 Table**). As a positive control, we also tagged a cell cycle-associated protein, ParB, and used CHA0 wild type without tagged proteins as a negative control. We collected the DNA after 3 h incubation with 9 µM of MMC. First, it is noteworthy that the addition of the V5 tag to the PrtR1 impacted the growth of the bacterium (**S4 Fig**). Indeed, the OD600nm dropped after 8 h suggesting that the stability of PrtR1 could be slightly impaired by the presence of the V5-tag, leading to the expression of the R-tailocin gene cluster and *in fine* R-tailocin release and cell lysis (**S4 Fig**). However, as we sampled at 3 h, we were unable to see this effect. Second, when induced, the PrtR1-V5 tagged strain showed higher OD600nm levels than the non-tagged wild type CHA0 (**S4 Fig**), which can be explained by the V5 impacting the functionality of the protein, rendering it less efficient.

As expected, we found that ParB binds regions near the origin and the terminus of replication (**S5 Fig**). Furthermore, there appears to be nonspecific binding or contaminant DNA in the regions with peaks for ribosomal RNA genes, as these peaks were observed in almost all conditions including in the CHA0 negative control but not in the ParB condition (**S6 Fig**). Focusing on PrtR1, we found that this regulator is highly specific to the promoter region of the R-tailocin gene cluster and not to any other viral-like cluster in CHA0 (**Fig 2B**).

Taken together, our results demonstrate that the LexA-like regulator PrtR1 binds directly to the promoter of the R-tailocin gene cluster of CHA0 and functions as a specific transcriptional repressor of this cluster as no crosstalk with the clusters encoding the other viral particles in the CHA0 genome was detected.

### MvaT and MvaV are global regulators that influence the expression of various competition traits including viral particles in CHA0

The binding of PrtR1 only partially explains the difference in expression of the R-tailocin gene cluster under non-induced and induced conditions. We therefore looked at other proteins that might be involved in the regulation of this gene cluster. We focused on the H-NS-like global regulators MvaT and MvaV in CHA0, as in *P. aeruginosa*, the homologs MvaT and MvaU function as master regulators of various virulence genes and as transcriptional silencers of xenogeneic DNA such as that from phages (29,34,38). The related MvaT and MvaV proteins of CHA0 were previously found to contribute to the control of the production of various exoproducts (30), but have not been investigated for the control of phage sequences in this strain so far.

To better understand which genes might be regulated by MvaT and MvaV when a cell is exposed to DNA damaging agents, we performed RNA-seq on a mutant lacking both of these genes (Δ*mvaT* Δ*mvaV*) under both non-inducing and inducing conditions (MMC and H2O2; **Fig 3A** and **Fig 3B**), along with mutants lacking either one or the other gene (Δ*mvaT* and Δ*mvaV*, **S7A Fig** and **S7B Fig**, respectively). Our results focus on the Δ*mvaT*Δ*mvaV* double mutant since *P. aeruginosa* homologs of MvaT and MvaV have been described to form heterodimers that act in concert to repress the expression of different genes (31). These proteins can also form homodimers that can still provide gene regulation and mask the effects of mutations to some extent (31). The RNA was harvested 3 h post induction to allow for comparison with results from the wild-type background (**S1C Fig**). To identify genes that are differentially regulated in the double mutant compared to the wild type under induction, we performed the following calculation: for each dataset (i.e., induction with H2O2 and MMC), we subtracted the differences observed between the double mutant Δ*mvaT*Δ*mvaV* and the CHA0 wild type in the non-induced condition from the differences observed in the induced condition **Fig 3A**, **3B**; for MMC: [Δ*mvaT*Δ*mvaV* vs WT]MMC - [Δ*mvaT*Δ*mvaV* vs WT]NI; for H2O2: [Δ*mvaT*Δ*mvaV* vs WT]H2O2 - [Δ*mvaT*Δ*mvaV* vs WT]NI). This approach allowed us to isolate the changes in gene expression that are specifically related to the double mutant when there is induction.

**Figure 3.**
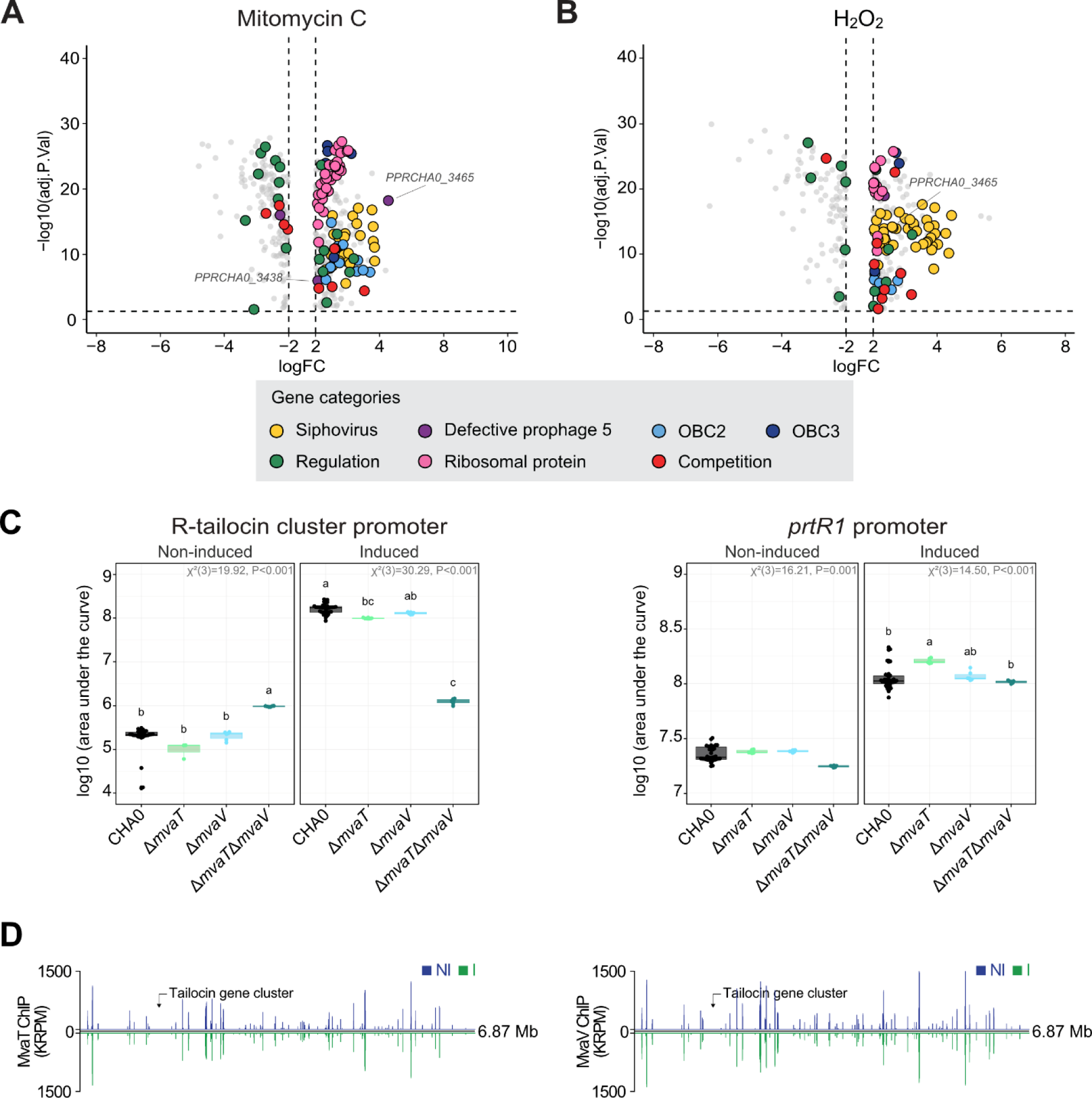
MvaT and MvaV are global regulators that influence the expression of many different competition traits including cell surface decorations, competition traits, and ribosomal proteins as well as the R-tailocin gene cluster of CHA0. (**A**, **B**) Volcano plots showing the RNA sequencing results of the effect of the Δ*mvaT*Δ*mvaV* deletion in CHA0 following exposure to 9 µM mitomycin C (MMC) (**A**) or 10 mM H2O2 (**B**). Genes that were differentially regulated in the double mutant compared to the wild type under induction, were identified using the following calculation: for each dataset (i.e., induction with MMC and H2O2), we subtracted the differences observed between the double mutant Δ*mvaT*Δ*mvaV* and the CHA0 wild type in the non-induced condition from the differences observed in the induced condition (for MMC: [Δ*mvaT*Δ*mvaV* vs WT]MMC - [Δ*mvaT*Δ*mvaV* vs WT]NI; for H2O2: [Δ*mvaT*Δ*mvaV* vs WT]H2O2 - [Δ*mvaT*Δ*mvaV* vs WT]NI). The colored dots correspond to different gene categories, yellow for the siphovirus, purple for the defective prophage 5, blue for the OBC2 and OBC3 clusters, green for regulation, pink for ribosomal proteins and red for competition. The expression of the R-tailocin cluster genes was not significantly different in the double mutant Δ*mvaT*Δ*mvaV* compared to the wild type in the induced condition. The vertical dashed lines correspond to the log2(Fold-change) thresholds set for this analysis (- 2 >log2(FC) > 2). The horizontal dashed line corresponds to the significance level of P < 0.05. (**C**) The expression of the R-tailocin gene cluster and the locus-specific regulatory gene *prtR1* was monitored using transcriptional reporters pOT1e-P*hol*-*egfp* and pOT1e-P*prtR1*-*egfp*, respectively, in the wild type CHA0 (black) and different *mvaT* and *mvaV* mutants (Δ*mvaT*, Δ*mvaV*, Δ*mvaT*Δ*mvaV*). Strains were grown in rich medium (NYB), following induction with 9 µM MMC or without induction. Optical density at 600 nm and GFP fluorescence (relative fluorescence units, RFU) were monitored every 10 min for 24 h in a BioTeK Synergy H1 plate reader. Detailed curves are shown in **S9 Figure**. The data were then used to calculate the area under the RFU curve. Statistical differences were assessed by Kruskal-Wallis tests with Bonferroni correction and are indicated by letters. (**D**) Chromatin immunoprecipitation sequencing (ChIP-seq) results of MvaT and MvaV under non-induced (NI, blue) and induced (I, 9 µM MMC, green) conditions.

Genes involved in bacterial competition traits, such as those encoding the CupB fimbriae, the siderophore pyoverdine, the antimicrobial metabolite DAPG, the Fit toxin, and a type VI secretion system (T6SS)-associated VgrG module (44,45) were upregulated under inducing conditions in Δ*mvaT*Δ*mvaV* (**Fig 3A**, **3B**). These findings support and extend previous findings pointing to a regulatory role of MvaT and MvaV in modulating competition traits of CHA0 (30). Furthermore, genes involved in cell wall / membrane envelope biogenesis (COG category M) which include lipopolysaccharide (LPS) cell surface decoration clusters, such as OBC2 and OBC3 (O-PS biosynthesis clusters), were also upregulated (**Fig 3A**, **3B**, **S8 Fig**) in this mutant, indicating that these regulators may influence bacterial interaction with its environment. Notably, the double mutant showed higher expression of genes related to translation (COG category J), including ribosomal protein genes, and exhibited some upregulation of genes involved in nucleotide metabolism and transport (COG category F) compared to the wild type, suggesting that MvaT and MvaV influence not only gene expression but also protein synthesis (**Fig 3A**, **3B**, **S8 Fig**).

In addition, some of the clusters encoding viral particles in CHA0 were also upregulated in Δ*mvaT*Δ*mvaV* under DNA-damaging conditions (**Fig 3A**, **3B**). Indeed, in both inducing conditions (i.e. MMC and H2O2), genes involved in the production and release of the *Siphoviridae* prophage were more upregulated compared to the wild type (**Fig 3A**, **3B**). Similarly to the wild type, the *Myoviridae* prophage was not upregulated in the Δ*mvaT*Δ*mvaV* mutant. Moreover, the double deletion resulted in the upregulation of two genes that were not differentially expressed in the wild type exposed to the DNA-damaging agents, PPRCHA0_3438 and PPRCHA0_3465, within a cluster that corresponds to the defective prophage 5 in the genome of *Pseudomonas protegens* Pf-5 (**Fig 3A**, **3B**) (46). Although the deletion of *mvaT* and *mvaV* affected the *Siphovirus* cluster of CHA0, the R-tailocin gene cluster was not significantly influenced in the Δ*mvaT*Δ*mvaV* mutant compared to the wild type CHA0 at the measured time point (3 h post induction), regardless of whether induction was performed with MMC or H2O2 (**Fig 3A**, **3B**).

As the RNA-seq analysis only provided a temporal snapshot of the induction process and could not capture dynamic changes, we examined the expression of the R-tailocin gene cluster and of *prtR1* in mutants lacking either *mvaT* (Δ*mvaT*), *mvaV* (Δ*mvaV*) or both genes (Δ*mvaT*Δ*mvaV*) over the course of 24 h (**Fig 3C**, **S9 Fig**). Although, the single mutation of either *mvaT* or *mvaV* did not affect the expression of the R-tailocin gene cluster or *prtR1* compared to the wild type, the deletion of both these genes affected the expression of the R-tailocin gene cluster of CHA0 (**Fig 3C**). Indeed, we found that, in non-inducing conditions the expression of the R-tailocin gene cluster in the Δ*mvaT*Δ*mvaV* mutant was significantly higher than in the wild type, while in inducing conditions it was significantly lower (**Fig 3C**). Interestingly, when comparing both non-induced and induced conditions, the expression of the R-tailocin gene cluster in the Δ*mvaT*Δ*mvaV* mutant did not differ (**Fig 3C**), suggesting that the MvaT and MvaV may acts as an indirect activator of the expression of the R-tailocin gene cluster. Furthermore, the expression levels of the negative regulatory gene *prtR1* in the mutants were similar as in the wild type in both non-induced and induced conditions (**Fig 3C**), suggesting that the effect of MvaT and MvaV on the expression of the R-tailocin gene cluster of CHA0 is independent of PrtR1. This result suggests that MvaT and MvaV may operate together to maintain R-tailocin expression at controlled levels even following exposure to DNA damaging agents.

To identify if MvaT and MvaV directly regulate the R-tailocin gene cluster of CHA0, we examined the regions that are bound by the two regulators using a ChIP-seq analysis. We confirmed that MvaT and MvaV are global regulators as they bind to more than 200 target sites throughout the CHA0 chromosome (**Fig 3D**). MvaT and MvaV also appear to bind to the same sites (**Fig 3D**, **S10 Fig**) supporting the fact that these proteins operate together as a heterodimer (31). As previously described for other H-NS-type regulators (29,38), MvaT and MvaV of CHA0 bind to AT-rich regions, i.e. regions with a significantly lower GC content than the average of the genome, which is at 63.4 % in the strain (47) (MvaT, *P* < 0.001; MvaV, *P* < 0.001, **S11A Fig**). Although the upstream region of *prtR1* is an AT-rich genomic region (58 %), MvaT and MvaV do not bind to it, nor to any sites upstream, downstream, or within the R-tailocin gene cluster (**S11B Fig**). Conversely, we found that MvaT and MvaV bind other cluster encoding viral particles (**S12 Fig**) and other genes involved in competition traits such as those involved in CupB fimbriae formation, T6SS VgrG modules, DAPG biosynthesis (**S13 Fig**) as well as clusters like OBC3 and OSA that are involved in O-antigenic LPS synthesis (39,48) (**S14 Fig**). Intriguingly, both regulators also bind to the promoter regions of their respective genes (**S15 Fig**), indicating a potential mechanism of autoregulation.

Taken together, these results demonstrate that MvaT and MvaV play a key role in regulating genes involved in bacterial competition traits, prophage activation, and cellular processes in CHA0. While they directly influence the expression of genes encoding the *Siphoviridae* prophage, T6SS components, and biosynthetic clusters for antimicrobial compounds by binding the clusters encoding these traits, they also indirectly regulate the expression of other clusters such as the one encoding the R-tailocins of CHA0.

## Discussion

The expression of phages and phage tail-like particles such as R-tailocins in bacteria needs to be tightly regulated as their production induces the lethal lysis of the producing cell. In this study, we set out to better identify how R-tailocin expression is regulated in environmental *Pseudomonas* using the model type strain CHA0.

### The SOS response is differentially regulated in the presence of MMC or H2O2

As expected, exposure to two known DNA-damaging agents, MMC and H2O2 (10), upregulated genes involved in the SOS response. In both conditions, *recN*, *cinA*, *recA*, *recX*, *sulA* and both *lexA* paralogs (*lexA* and *lexA2*) were upregulated (**Fig 1A**, **1B**). However, in the presence of MMC, *dnaE2*, *dinB* and *uvrA_2* were additionally upregulated compared to the H2O2 condition (**Fig 1A**, **1B**). This differential expression of the genes depending on the condition could be due to a different response depending on the stress induced and the time scale. Genes involved in the SOS response are expressed sequentially, and not all are induced at the same level (14). Indeed, the SOS response is precisely regulated and expressed depending on the damage inflicted (14). Accordingly, although the R-tailocin genes were highly upregulated when in the presence of both compounds, genes involved in the siphovirus production were slightly less upregulated when CHA0 was in the presence of H2O2 compared to MMC (**Fig 1A**, **1B**). Conversely, the genes associated with the *Myoviridae* prophage were not differentially expressed in the two conditions (**Fig 1A**, **1B**). This is in line with previous findings, as the myovirus is considered a cryptic prophage and is not produced when cells are induced with either compound (1).

Genes involved in the SOS system could be regulated by other stress responses. In *E. coli* the alternative sigma factor RpoS, which regulates the response to nutrient starvation during the stationary phase (49), as well as the primary sigma factor RpoH, which is essential for cell viability (50), were shown to induce the expression of some of the SOS system-related genes (14). Furthermore, H2O2 is known to induce the oxidative stress response via OxyR and has been shown to potentially interact with some SOS genes such as *recN* (51). Consequently, tailocin and phage expression may be induced by different stress responses and may allow bacterial populations to maintain low levels of tailocin expression even in the absence of DNA-damaging agents.

### PrtR1 tightly controls R-tailocin expression in CHA0, acting as its primary transcriptional repressor

The regulation of tailocin gene expression in response to DNA damage has been extensively studied in the opportunistic human pathogen *P. aeruginosa*. In this bacterium, PrtR acts as a negative regulator, analogous to PrtR1 in CHA0, but it also relies on PrtN, a transcriptional activator of the tailocin gene cluster, which is absent in CHA0 (1,20). In *P. aeruginosa*, PrtR represses *prtN* expression under non-inducing conditions (20). Upon exposure to DNA-damaging agents, nucleoprotein filaments of RecA activate the autoproteolytic cleavage of PrtR, resulting in the deblocking of PrtN (15), which in turn activates the expression of the tailocin genes by binding to the promoter of the cluster (20). Here we demonstrate that PrtR1 directly regulates the expression of the R-tailocin gene cluster in the model environmental pseudomonad CHA0 (**Fig 2**). Deletion of *prtR1* leads to upregulation of the tailocin gene cluster under both induced and non-induced conditions (**Fig 2A**), demonstrating that PrtR1 serves as a direct inhibitor of R-tailocin production similarly to LexA that following exposure to DNA-damaging agents is cleaved, permitting the expression of SOS response genes (14,15). ChIP-seq analysis revealed that PrtR1 binds to the promoter of the R-tailocin gene cluster (**Fig 2B**). Notably, PrtR1 binding appeared to be stronger in non-induced conditions compared to induced conditions, further supporting its role as a specific repressor of R-tailocin expression in CHA0.

Interestingly, in *Pseudomonas fluorescens* SF4c, a PrtR-type protein with 91 % identity to PrtR1 in CHA0 and identical predicted functional domains has been described as a positive rather than a negative regulator of the R-tailocin gene cluster (42). We identified an operator-like sequence (ATAAATGCATTTAC) previously described in *P. fluorescens* SF4c (42) within the promoter region of the R-tailocin gene cluster in CHA0, at the ChIP-seq peak with the highest signal intensity. However, in CHA0, this sequence is located 391 bp upstream of the *hol* gene, whereas in *P. fluorescens* SF4c, it is positioned closer to the start site of the gene (42). While the operator sequence of the PrtR orthologs is conserved between CHA0 and *P. fluorescens* SF4c, and the orthologs share a high percentage of identity, they could have different functions in these two strains.

Two additional PrtR-type proteins are encoded in the genome of CHA0, i.e. PrtR2 (PPRCHA0_2149) associated with a defective bacteriocin-encoding prophage and PrtR3 (PPRCHA0_3804) associated with the siphovirus prophage. PrtR1 binds to the promoter region of the R-tailocin gene cluster, and the expression of this cluster was not affected in mutants lacking the two other phage-like clusters (Δ*myo*Δ*siph*; **Fig 2A**). Based on the observations, we assume that the gene clusters encoding phage tail-like particles and phages in CHA0 have specialized and non-interchangeable locus-specific regulators. Interestingly, the expression of *prtR1* was affected not only in the mutant lacking this gene (Δ*prtR1\**) but also in the mutant lacking the entire R-tailocin gene cluster (Δtailcluster) (**Fig 2A**). As the Δ*prtR1\** mutant is a “conditional” mutant that lacks both the entire R-tailocin gene cluster and *prtR1*, we hypothesized that a gene within the cluster may play a role in regulating *prtR1*. To test this hypothesis, we used transcriptional reporters for the R-tailocin gene cluster and *prtR1* in mutants lacking specific genes within the R-tailocin gene cluster that we suspected to encode putative regulators (Δ*lateD*, Δ*PPRCHA0_1250*, Δ*PPRCHA0_1251* and Δ*PPRCHA0_1252*). However, we did not observe any notable differences in the expression of the cluster (**S3 Fig**).

### MvaT and MvaV indirectly regulate R-tailocin expression in CHA0 but directly regulate some other competition traits

Besides PrtR1, additional regulators are likely involved in the control of R-tailocin expression. We focused on MvaT and MvaV, which belong to a family of H-NS-like global regulators previously described to be involved in the expression of virulence factors and prophages in *P. aeruginosa* (29,34,38). Interestingly, while the related *mvaT* and *mvaU* genes in *P. aeruginosa* PAO1 are downregulated in response to ciprofloxacin (13), our RNA-seq analysis showed that *mvaT* and *mvaV* in CHA0 were not differentially expressed upon exposure of the bacterium to MMC or H2O2 (**Fig 1A**, **1B**). However, we found that MvaT and MvaV have an indirect regulatory effect on the expression of the CHA0 R-tailocin gene cluster (**Fig 3**). In the Δ*mvaT*Δ*mvaV* double mutant, the R-tailocin cluster showed comparable expression levels under non-inducing and inducing conditions, in contrast to the wild type where induction led to a significant upregulation (**Fig 3C**). Despite the clear effect on R-tailocin cluster expression in CHA0, MvaT and MvaV did not bind to its promoter region (**S11B Fig**). We propose two different mechanisms by which the MvaT-MvaV heterodimer may regulate the expression of the R-tailocin gene cluster in CHA0. As H-N-type regulators have been described as silencers (28,29), we suggest that MvaT and MvaV may either repress a repressor or repress an activator. First, acting as an anti-repressor, MvaT and MvaV would lead to the activation of the expression of the R-tailocin cluster in a wild-type background. In a mutant lacking both *mvaT* and *mvaV* there would be an inability to fully activate the expression of the cluster in induced conditions. However, MvaT and MvaV do not directly regulate the expression of the negative regulator PrtR1, as they do not bind to the *prtR1* promoter region (**S11B Fig**), and the expression of *prtR1* is not significantly altered in the Δ*mvaT*Δ*mvaV* mutant (**Fig 3C**). Thus, the most likely hypothesis is the second, where MvaT and MvaV would act as an anti-activator, leading to the repression of the R-tailocin gene cluster in a wild type background. In an Δ*mvaT*Δ*mvaV* mutant, there would be increased activation of the R-tailocin gene cluster in a non-induced condition (**Fig 3C**).

Similar to their homologs MvaT and MvaU in *P. aeruginosa* (29,33), MvaT and MvaV were confirmed as global regulators binding to more than 200 chromosomal targets in CHA0 (**Fig 3D**). Thus, this regulation of the R-tailocin gene cluster of CHA0 could involve many different intermediates. As H-NS-type proteins, MvaT and MvaV typically target AT-rich regions, permitting the selective silencing of xenogenic elements, such as genes acquired by horizontal gene transfer, prophage sequences and pathogenicity islands, which are often lower in GC content than the rest of the bacterial genome (24,27,29,33,34,52). This AT-rich binding preference was also observed for MvaT and MvaV in CHA0 (**S11A Fig**).

In *P. aeruginosa*, MvaT and MvaU co-regulate the expression of over 100 genes (29), including genes for pyocyanin production, biofilm formation, type III secretion, and regulatory elements such as RpoS (29,31,34). Similarly, MvaT and MvaV in CHA0 influence diverse competition traits, including antimicrobial production, motility, cell surface properties and biocontrol activity (30). Our results support these observations as in a mutant lacking both regulators (Δ*mvaT*Δ*mvaV*), genes involved in the formation of competitive traits such as CupB fimbriae, a T6SS-related VgrG module, the siderophore pyoverdine, the antimicrobial DAPG, and the toxin Fit were upregulated under inducing conditions (**Fig 3A**)(44,45). Additionally, gene clusters specifying cell surface decorations, such as the LPS O-antigens OBC2 and OBC3, were also upregulated (**Fig 3A**). Notably, not all these genes had upstream binding sites for MvaT and MvaV, suggesting that MvaT and MvaV may be directly and indirectly involved in the modulation of interactions of *P. protegens* with the environment, including plant and insect hosts.

## Conclusion

To conclude, we propose a model for the regulation of the R-tailocin gene cluster in *Pseudomonas protegens* CHA0, involving PrtR1, MvaT, and MvaV (**Fig. 4**). Under non-inducing conditions, PrtR1 directly represses the expression of the R-tailocin gene cluster, with its own expression being influenced by a protein within the R-tailocin cluster (**Fig. 4A**). Under inducing conditions, the SOS system is upregulated in response to stress. RecA* nucleoprotein filaments can then induce the autocatalytic cleavage of PrtR1 (**Fig. 4B**). This relieves the repression from the promoter of R-tailocin gene cluster, permitting the expression of the R-tailocin gene cluster, and consequently the formation and release of R-tailocin particles from CHA0 cells (**Fig. 4B**). In both non-inducing and inducing conditions, MvaT and MvaV, while not directly binding to the R-tailocin gene cluster, indirectly influence its expression (**Fig. 4**). Other intermediates involved in the regulation of these viral-like particles remain to be discovered in order to fully understand how their production is controlled. Overall, this study provides new insights into the complex regulatory networks governing the production of antimicrobial molecules in CHA0, highlighting the dynamic interplay between phage tail-like particles, stress responses, global gene regulators, and secondary metabolite expression.

**Figure 4.**
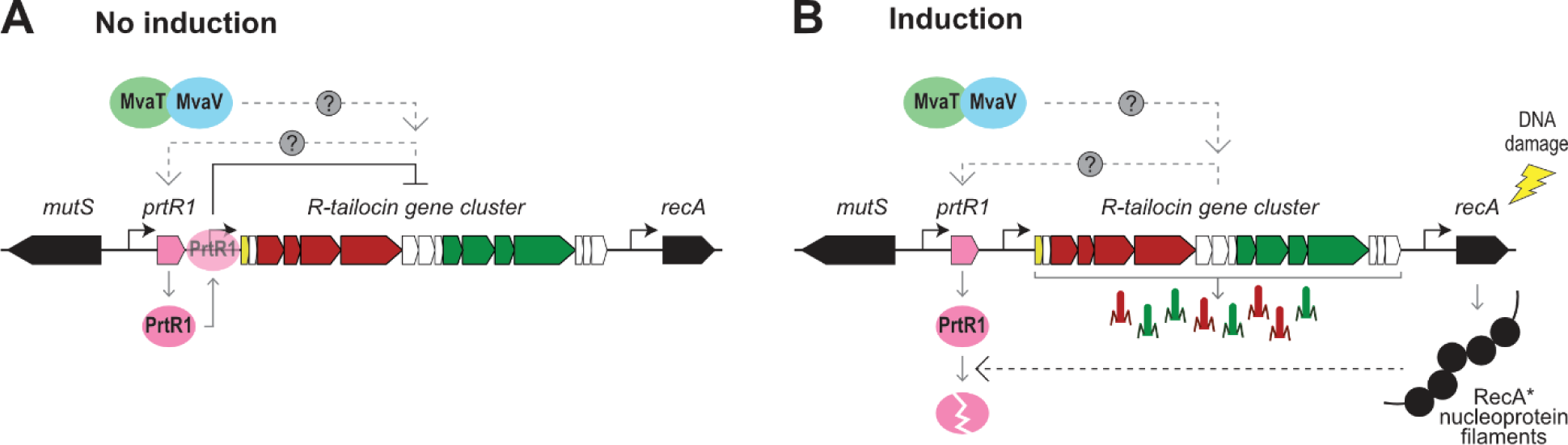
Proposed model for the regulation of the expression of the R-tailocin gene cluster of *P. protegens* CHA0 under non-induced conditions (A) and upon induction with a DNA-damaging agent (B). (**A**) In the non-induced condition, the expression of the R-tailocin gene cluster is directly inhibited by the binding of the LexA-related repressor PtrR1 to the promoter region of the cluster. MvaT and MvaV are indirectly involved in the expression of the R-tailocin gene cluster. The expression of PrtR1 is possibly regulated by other genes within the R-tailocin gene cluster itself. (**B**) Under inducing conditions, danger sensing systems such as the bacterial SOS response are activated. Activated RecA*, bound to damaged single-stranded DNA will induce the autocatalytic cleavage of PrtR1, relieving the promoter region of the R-tailocin gene cluster. As in a non-induced condition, MvaT and MvaV also appear to be indirectly involved in the expression of the R-tailocin gene cluster. The genes encoding the structural parts of the R-tailocin #1 and the R-tailocin #2 are colored red and green, respectively, the *prtR1* gene is colored pink, the gene encoding the lytic gene holin (*hol*) is colored yellow, other genes belonging to the cluster are colored white, and the bacterial genes neighboring the R-tailocin gene cluster are colored black. MvaT and MvaV proteins are colored in green and blue, respectively.

## Material and methods

### Bacterial strains, plasmids, media, and culture conditions

All plasmids and strains used in this study are listed in **S1** and **S2 Tables**, respectively. *Pseudomonas* strains were routinely cultured on nutrient agar (NA) and in nutrient yeast broth (NYB) at 25 °C. *Escherichia coli* was grown in lysogeny broth (LB) at 37 °C. Appropriate antibiotics (kanamycin, 25 µg mL^-1^; gentamicin, 10 µg mL^-1^) were used when required.

### Mutant construction

Mutants of CHA0 were constructed as in Heiman *et al.*, 2022 (39), using the suicide vector pEMG and the I-SceI system (53) with a protocol adapted to CHA0 (1). Plasmids and primers are listed in **S1** and **S3 Tables**. The mutants are listed in **S2 Table**.

### Transcriptional reporter construction

To construct the different transcriptional reporters used in this study, the promoter regions of the gene / gene cluster of interest were amplified by PCR and cloned upstream of the *egfp* gene into the expression vector pOT1e (47). To construct the transcriptional reporter for the R-tailocin gene cluster (pOT1e-P*hol*-*egfp*), a 516-bp DNA region upstream of the holin gene (*hol*) (intergenic region between the *prtR1* and the *hol* genes, PPRCHA0_1217 and PPRCHA0_1218, respectively) was amplified (**Fig 2A**). To construct the transcriptional reporter of the predicted negative regulator PrtR1 of the R-tailocin gene cluster (pOT1e-P*prtR1*-*egfp*), a 50-bp sequence in the intergenic region upstream of *prtR1* was amplified (**Fig 2A**). The resulting plasmids (**S1 Table**), as well as an empty version of the pOT1e vector, were transformed by electroporation into wild type CHA0 and mutant derivatives (Δ*prtR1\**, Δtailcluster, Δ*myo*Δ*siph*, Δ*mvaT*Δ*mvaV*, Δ*mvaT*, Δ*mvaV*, Δ*lateD*, Δ*PPRCHA0_1250*, Δ*PPRCHA0_1251* and Δ*PPRCHA0_1252*). Transformants were selected on NA plates supplemented with gentamicin.

### RNA extraction

An RNA sequencing (RNA-seq) approach was used with either mitomycin C (MMC) or H2O2, two DNA-damaging agents known to induce the SOS response and consequently the expression of the R-tailocin gene cluster (8,10,15). To identify the appropriate concentration of H2O2 for induction, five different concentrations (100 mM, 10 mM, 0.1 mM, 10 μM and 0.1 μM) were first tested using the R-tailocin gene expression reporter strain CHA0 pOT1e-P*hol*-*egfp* (**S1A**, **S1B Fig**). Briefly, an overnight culture of CHA0 pOT1e-P*hol*-*egfp* was restarted (1:100) in fresh NYB. When the culture reached the exponential growth phase (OD600nm of 0.4-0.6), the OD600nm was adjusted to 1.0 in fresh NYB. Twenty µL of the adjusted bacterial culture were added to 180 µL of NYB supplemented with H2O2 at the different concentrations tested. GFP fluorescence (excitation 479 nm, emission 520 nm) and OD600nm were monitored for 24 h with time points every 10 min using a BioTek Synergy H1 plate reader (BioTek Instruments Inc., Winooski, VT, USA). Ten mM of H2O2 was the best condition for inducing the R-tailocin gene expression reporter (**S1A**, **S1B Fig**).

The RNA-seq approach was performed on the wild type CHA0 and mutants defective for the H-NS-like regulators MvaT and MvaV (Δ*mvaT*, Δ*mvaV* and Δ*mvaT*Δ*mvaV*). Overnight cultures of the strains were restarted into fresh NYB at a ratio of 1:100. When the bacterial cultures reached the exponential growth phase (OD600nm of 0.4-0.6), they were either induced with 9 µM MMC or 10 mM H2O2 or were not induced. Following a 3 h induction, cells were treated with RNAprotect Bacteria Reagent (Qiagen) for 5 min at room temperature according to the manufacturer’s instructions. Cultures without the inducer were used as a control. All the sample were centrifuged at 5000 *g* for 10 min and the pellets were flash-frozen in liquid nitrogen and stored at -80 °C prior to further processing. RNA was extracted using the RNeasy Mini Kit (Qiagen). Potential DNA contamination was removed by TurboDNase treatment (ThermoScientific), and the RNA was purified using the RNAeasy kit (Qiagen). Aliquots of the three replicates from the 12 conditions were sampled to perform CFU counting prior to induction and 3 h post induction.

### RNA-seq DNA library preparation and Illumina sequencing

Ribosomal RNA was removed using the RiboCop rRNA depletion kit (Lexogen). The RNA-seq libraries were prepared using the Illumina Stranded mRNA kit (Illumina) according to the manufacturer’s recommendations. The final library was purified with SPRI beads at a 0.9X ratio and quantified using Qubit (Thermo Scientifics). The size pattern of the final library was analyzed with a fragment analyzer (Agilent). Single-end sequencing was performed on a NovaSeq6000 system (Illumina), 100 cycles. Sequencing data were demultiplexed using the bcl2fastq Conversion Software v2.20 (Illumina) and further processed for differential expression analysis.

### RNA-seq sequence processing and statistical analysis

The adapters of the purity-filtered reads were trimmed and poor quality reads were removed with Cutadapt v. 2.5 (54). Reads matching to ribosomal RNA sequences were removed with Fastq_screen (v. 0.11.1). The remaining reads were further filtered for low complexity with Reaper v. 15-065 (55). Reads were aligned to the CHA0 genome (accession number: LS999205) using STAR v. 2.5.3a (56). The number of read counts per gene locus was summarized with htseq-count v. 0.9.1 (57) using the CHA0 gene annotation. The quality of the RNA-seq data alignment was assessed using RSeQC v. 2.3.7 (58). Statistical analysis was performed in R (R version 4.2.2). Genes with low counts were filtered out according to the rule of 1 count per million (cpm). TMM normalization was used to scale library sizes. The normalized counts were then transformed to cpm values and a log2 transformation was applied by means of the function cpm with the parameter setting prior.counts = 1 (EdgeR v 3.30.3; (59)). The characteristics of the RNA-seq are described in **S4 Table**.

### Transcriptional reporter expression monitoring

The expression of the transcriptional reporters of the gene / gene cluster of interest (pOT1e-P*hol*-*egfp* and pOT1e-P*prtR1*-*egfp*) was monitored in different mutant backgrounds (CHA0 wild type, Δ*myo*Δ*siph*, Δ*prtR1\**, Δtailcluster, Δ*mvaT*Δ*mvaV*, Δ*mvaT*, Δ*mvaV*, Δ*lateD*, Δ*PPRCHA0_1250*, Δ*PPRCHA0_1251* and Δ*PPRCHA0_1252*). To this end, overnight cultures were restarted into fresh NYB at a ratio of 1:100. When the cultures reached the exponential growth phase (OD600nm of 0.4-0.6), they were inoculated into 96-well plates prepared with 180 μL per well of NYB, supplemented or not with 9 µM MMC, adjusting the starting OD600nm in the wells to 0.1. The OD600nm and the GFP fluorescence were monitored every 10 min for 24 h using a BioTek Synergy H1 plate reader (BioTek Instruments Inc., Winooski, VT, USA). A minimum of three biological replicates with each three technical replicates were performed. Data were analyzed and figures were drawn using R studio version 4.3.1.

### Chromatin immunoprecipitation (ChIP)

To identify the binding regions of putative R-tailocin gene cluster-regulating proteins, a chromatin immunoprecipitation sequencing (ChIP-seq) analysis was performed. We focused on four proteins: the predicted repressor PrtR1 of the R-tailocin gene cluster, the two H-NS-like global regulators MvaT and MvaV, and the cell cycle-associated protein ParB as a control. Plasmids containing each gene encoding the protein of interest flagged with a V5 in C-terminal (synthesized by GenScript) were cloned into the suicide vector pEMG (**S1 Table**) to construct the V5-tagged CHA0 derivatives *prtR1*-V5, *mvaT*-V5, *mvaV*-V5, and *parB*-V5 (**S2 Table**) for the ChIP-seq. V5-tagged derivative strains were compared with the wild type CHA0 for differences in growth by monitoring the OD600nm every 10 min for 24 h using a BioTek Synergy H1 plate reader as described above. The time point at which the cultures were harvested for ChIP was determined by immunoblot analysis (**S16 Fig**). Overnight cultures of the strains were restarted at a 1:100 ratio in fresh NYB. When the cultures reached the exponential growth phase (OD600nm=0.4-0.6), 9 µM MMC was added. Samples were collected prior to induction and 1 h, 2 h and 3 h post induction and immunoblot analysis was performed according to the protocol of Chai *et al.* (60) using anti-V5 (1:2000) as the first antibody and anti-mouse (1:150000) as the second antibody. Three hours post induction was chosen as the best time point to collect samples for ChIP.

Samples from cultures of the different strains (*mvaT*-V5, *mvaV*-V5, *prtR1*-V5 and *parB*-V5, CHA0 wild type as a negative control) grown under the conditions described above were collected 3 h post induction for ChIP analysis following a protocol adapted for CHA0 from Antar *et al*. (61,62). Briefly, 1 mL of fixation buffer F [50 mM tris-HCl, pH 7.4, 100 mM NaCl, 0.5 mM EGTA, pH 8.0, 1 mM EDTA, pH 8.0, and 10 %, w/v formaldehyde] was added to the 10 mL of culture and incubated for 30 min at room temperature. Cells were harvested by centrifugation (5000 *g* for 10 min at 4 °C) and washed with 1x PBS. Samples adjusted for an OD600nm of 1 in 2 mL were resuspended in TSEMS lysis buffer [50 mM tris pH 7.4, 50 mM NaCl, 10 mM EDTA pH 8.0, 0.5 M sucrose and PIC (Sigma-Aldrich), and 6 mg mL^-1^ lysozyme from chicken egg white (Sigma-Aldrich)] and incubated for 30 min at 37 °C. Protoplasts were then washed twice with TSEMS lysis buffer. Supernatants were discarded and pellets were flash-frozen and stored at -80 °C until further use. Anti V5 was preincubated with Protein G–coupled dynabeads (Invitrogen) in a 1:1 ratio for 2 h at 4 °C on a rotating wheel. Beads were then washed with buffer L [50 mM Hepes-KOH pH 7.5, 140 mM NaCl, 1 mM EDTA pH 8.0, 1 % (v/v) Triton X-100, 0.1 % (w/v) sodium deoxycholate, ribonuclease A (0.1 mg mL^-1^), and PIC (Sigma-Aldrich)], aliquoted into each sample and incubated for 2 h at 4 °C. A series of washes were then performed on the beads conjugated with the proteins with buffer L, buffer L5 [buffer L containing 500 mM NaCl], buffer W [10 mM Tris-HCl pH 8.0, 250 mM LiCl, 0.5 % (v/v) NP-40, 0.5 % (w/v) Na deoxycholate, and 1 mM EDTA pH 8.0], and buffer TE [10 mM Tris-HCl pH 8.0, and 1 mM EDTA pH 8.0]. Beads were resuspended in 520 μL TES buffer [50 mM Tris-HCl pH 8.0, 10 mM EDTA pH 8.0, and 1 % (w/v) SDS]. Samples were incubated overnight at 65 °C with shaking to remove the DNA from the beads. The DNA was then purified using a phenol-chloroform extraction. Five hundred μL of phenol was added to the samples and mixed by vortexing. Samples were centrifuged at 20000 *g* for 10 min, and 450 μL of the aqueous phase was transferred to a new tube and mixed with an equal volume of chloroform. Samples were centrifuged before recovering 400 μL of the aqueous phase. DNA was precipitated with 1 mL of 100 % ethanol (2.5 × volume), 40 μL of 3 M NaOAc (0.1 × volume), and 1.2 μL of GlycoBlue and incubated for 20 min at -20 °C. Samples were centrifuged at 20,000 *g* for 10 min. The resulting pellets were purified with a PCR purification kit (Qiagen) and eluted in 50 μL EB buffer.

### ChIP DNA library preparation and Illumina sequencing

ChIP DNA samples were quantified using Qubit (Thermo Scientifics) to determine the quantity of DNA to use. Depending on the concentration measured by the Qubit, between 0.35-2 ng of DNA were used for sequencing. The DNA was sheared using a Covaris S220 (settings: 50 µL in microTUBES with AFA fiber; peak incident power, 175 W; duty factor, 10 %; cycles per burst, 200; treatment time, 120 s) and purified with SPRI beads at a 2.0 X ratio. The library was prepared using the NuGen Ovation Ultra Low system v2 (Tecan Trading AG) with 15 cycles of PCR amplification and using a unique dual indexing strategy. The final libraries were quantified using Qubit and their quality assessed on a Fragment Analyzer. Sequencing was performed on an Illumina NovaSeq 6000 for 300 cycles (paired end 150 nt reads). Sequencing data were demultiplexed using the bcl2fastq2 Conversion Software v2.20 (Illumina).

### ChIP-seq sequence processing and statistical analysis

Unique molecular identifiers (UMI) were added in the reads for both R1 and R2 paired reads fastq files using an in-house Perl script. Adapters were trimmed using trim_galore.0.6.4 (Martin, 2011) and both raw fastq and trimmed fastq reads were quality-checked using fastQC version 0.11.9 (https://www.bioinformatics.babraham.ac.uk/projects/fastqc/). The trimmed reads were mapped to the genome of CHA0 (accession number LS999205) using Bowtie version 2.4.4 with a minimum fragment length of 10, a maximum fragment length of 700, with discordant alignments for paired reads and unpaired alignments suppressed. Duplicate reads were removed using Picard tools 2.5.0 (https://broadinstitute.github.io/picard/). Bigwig files were generated using bamCoverage from the DeepTools version 3.5.1 (63). These files were used to visualize the results with IGV and to generate the figures. Bed files in BEDPE format, generated from the bam files using bedtools bamtobed with final fragment positions using a local Perl script, were used for peak calling with MACS (macs2 callpeak with the narrow peak option, FDR set to 0.05, genome size set to 7 M). Plots were generated using R version 4.3.1 and IGV version 2.16.2. The characteristics of the ChIP-seq are described in **S5 Table**.

### Statistics and reproducibility

The number of biological and technical replicates performed for each experiment is detailed in the figure legends. Data were analyzed using R studio version 4.3.1 and considered significantly different at *P* < 0.05. Data were tested for normal distribution and homogeneity of variance and transformed using the Shapiro-Wilk and Bartlett tests, respectively. ANOVA coupled with HSD-Tukey test was performed. When the normal distribution was not respected, non-parametric tests such as Wilcoxon or Kruskal-Wallis tests with Bonferroni correction were used to assess significant differences between conditions.

## Data availability

The raw data of the RNA-seq and the ChIP-seq are available from ArrayExpress under accession numbers E-MTAB-14715 and E-MTAB-14721, respectively. All other data are available on Zenodo (https://doi.org/10.5281/zenodo.14072069).

## Acknowledgements

We thank the Genomic Technologies Facility (GTF) at the University of Lausanne, specifically Viviane Praz, Corinne Peter, Johann Weber and Julien Marquis for their help with library preparation, sequencing and analysis of ChIP-seq data. We also thank Stephan Gruber for his insightful comments and corrections on the manuscript.

## Supporting information captions

**S1 Fig. H2O2 and mitomycin C induce the expression of the R-tailocin gene cluster and impact the growth of CHA0 wild type and H-NS mutants.** (**A**, **B**) R-tailocin gene expression was monitored using *P. protegens* CHA0 carrying a transcriptional reporter vector with a copy of the R-tailocin gene cluster promoter located upstream of the *egfp* gene. Optical density at 600 nm (**A**, OD600nm) and GFP fluorescence (**B**, relative fluorescence units, RFU) were monitored in rich medium (NYB) following induction with different concentrations of H2O2 or 9 µM mitomycin C by taking measurements every 10 min in a BioTeK Synergy H1 plate reader. The curves with error bars represent the average (± standard deviation) of measurements from three biological replicates with three technical replicates each. The red bar corresponds to the three-hour time point at which samples were collected for RNA sequencing (RNA-seq). (**C**) Colony forming units (CFU) of the different samples used for the RNA-seq for *P. protegens* CHA0 wild type, Δ*mvaT*, Δ*mvaV* and Δ*mvaT*Δ*mvaV*. Samples were collected prior to induction (at time 0 h) and 3 h after induction with H2O2 (10 mM), mitomycin C (9 µM) or without induction (control). Three technical replicates were collected for each condition.

**S2 Fig. Growth (OD600nm) and relative fluorescence (RFU) curves of transcriptional reporters of R-tailocin gene cluster and *prtR1* expression in CHA0 wild type and mutants Δ*prtR1\**, Δtailcluster, and Δ*myo*Δ*siph*.** The expression of the R-tailocin gene cluster and the locus-specific regulatory gene *prtR1* was monitored using the transcriptional reporters pOT1e-*Phol-egfp* and pOT1e-P*prtR1-egfp*, respectively, in the wild type CHA0 (black) and its mutants Δ*prtR1\** (pink), Δtailcluster (burgundy) and Δ*myo*Δ*siph* (purple). Strains were grown in rich medium (NYB), following induction with 9 µM mitomycin C or without induction. Optical density at 600 nm (OD600nm) and GFP fluorescence (relative fluorescence units, RFU) were monitored in rich medium (NYB) every 10 min for 24 h in a BioTeK Synergy H1 plate reader. Curves show means (± standard deviation) of a minimum of three biological replicates with three technical replicates each.

**S3 Fig. Effect of the deletion of genes with suspected regulatory function on the expression of the R-tailocin gene cluster and *prtR1* in CHA0.** The expression was monitored using the transcriptional reporters pOT1e-*Phol-egfp* and pOT1e-P*prtR1-egfp*, respectively, in the wild type CHA0 (black) and its different mutants (Δ*prtR1\**, Δtailcluster, Δ*lateD*, Δ*PPRCHA0_1250*, Δ*PPRCHA0_1251* and Δ*PPRCHA0_1252*). Strains were grown in rich medium (NYB), following induction with 9 µM mitomycin C or without induction. (**A**) Optical density at 600 nm and GFP fluorescence (relative fluorescence units, RFU) were monitored every 10 min for 24 h in a BioTeK Synergy H1 plate reader. (**B**) The data were then used to calculate the area under the RFU curve. Statistical differences were assessed by Kruskal-Wallis tests with Bonferroni correction and are indicated by letters. A minimum of three biological replicates with at least three technical replicates are plotted. Curves show means (± standard deviation) of a minimum of three biological replicates with three technical replicates each.

**S4 Fig. Growth kinetics of CHA0 derivatives expressing the different V5-tagged proteins (MvaT-V5, MavV-V5, PrtR1-V5 and ParB-V5).** To test for differences in growth patterns caused by the V5-flagged proteins, the growth of the different strains including the wild type CHA0 in rich medium (NYB) under non-induced and induced (9 µM mitomycin C) conditions was monitored for 24 h by measuring the optical density at 600 nm (OD600nm) every 10 min in a BioTeK Synergy H1 plate reader. Curves show means (± standard deviation) of two biological replicates with four technical replicates each. The red bar in the lower panels, representing the first five hours of incubation, corresponds to the three-hour time point at which samples were collected for RNA sequencing.

**S5 Fig. ParB binds specifically to regions near the origin and terminus of replication of CHA0.** Chromatin immunoprecipitation sequencing (ChIP-seq) results of ParB-V5 over the origin and terminus of replication under non-induced (NI, blue) and induced (I, 9 µM mitomycin C, green) conditions. The genes involved in replication or DNA repair are colored orange and genes involved in transport are colored brown. Other genes are colored black.

**S6 Fig. DNA encoding ribosomal RNA is a highly abundant contaminant in all conditions of the ChIP-seq including the condition without V5-flagged proteins (CHA0 wild type).** Chromatin immunoprecipitation sequencing (ChIP-seq) for *P. protegens* CHA0 wild type, MvaT-V5, MvaV-V5, PrtR1-V5 and ParB-V5 and with highlighted results of the five different clusters (annotated from 1 to 5) encoding ribosomal RNA in *P. protegens* CHA0, in non-induced (NI, blue) and induced (I, green, 9 µM mitomycin C) conditions. Each cluster encodes at least one 16S, one 23S and one 5S ribosomal subunit. The genes encoding ribosomal RNA and transfer RNA (tRNA) are colored light blue and dark blue, respectively, and other genes are colored black.

**S7 Fig. The deletion of *mvaT* or *mvaV* leads to the differential regulation of prophage clusters as well as gene clusters related to cell surface decorations and competition traits in CHA0.** Volcano plots showing the RNA sequencing results of the effect of the deletion of *mvaT* (**A**) and *mvaV* (**B**) in *P. protegens* CHA0 following exposure to 9 µM mitomycin C (left) or 10 mM H2O2 (right). The colored dots correspond to different gene categories, yellow for the siphovirus, purple for the defective prophage 5, blue for the OBC2, OBC3 and OSA clusters, green for regulation, pink for ribosomal proteins and red for competition. The vertical dashed lines correspond to the log2(Fold-change) thresholds set for this analysis (-2 >log2(FC) > 2). The horizontal dashed line corresponds to the significance level of *P* < 0.05.

**S8 Fig. The deletion of both *mvaT* and *mvaV* leads to a higher expression of genes related to translation compared to the CHA0 wild type.** Genes that are down-regulated and up-regulated in the Δ*mvaT*Δ*mvaV* mutant compared to the CHA0 wild type according to their COG assignment following exposure to the two compounds. The length of the bars corresponds to the percentage of genes associated with the different COG assignments.

**S9 Fig. Growth (OD600nm) and relative fluorescence (RFU) curves of transcriptional reporters of R-tailocin gene cluster and *prtR1* expression in CHA0 wild type, single mutants Δ*mvaT*, Δ*mvaV* and double mutant Δ*mvaT*Δ*mvaV*.** The expression of the R-tailocin gene cluster and the locus-specific regulatory gene *prtR1* was monitored using the transcriptional reporters pOT1e-*Phol-egfp* and pOT1e-P*prtR1-egfp*, respectively, in the wild type CHA0 (black) and mutants Δ*mvaT*, Δ*mvaV*, Δ*mvaT*Δ*mvaV*. Strains were grown in rich medium (NYB), following induction with 9 µM mitomycin C or without induction. Optical density at 600 nm (OD600nm) and GFP fluorescence (relative fluorescence units, RFU) were monitored in rich medium (NYB) every 10 min for 24 h in a BioTeK Synergy H1 plate reader. Curves show means (± standard deviation) of a minimum of three biological replicates with two technical replicates each.

**S10 Fig. Comparison of the ChIP-seq peaks of MvaT and MvaV under non-induced (A) and induced conditions (B).** Chromatin immunoprecipitation sequencing (ChIP-seq) results of MvaT-V5 (green highlight) and MvaV-V5 (blue highlight) under non-induced (blue peaks) and induced (9 µM mitomycin C, green peaks) conditions.

**S11 Fig. MvaT and MvaV of CHA0 are global regulators preferentially binding regions with a high A-T percentage but not the R-tailocin gene cluster of CHA0.** (**A**) GC content of the reads sequenced under the different peaks of MvaT and MvaV. The red vertical line corresponds to the average GC content of the *P. protegens* CHA0 genome (63.4 %). Statistical differences were assessed by Kruskal-Wallis tests coupled with Dunn tests, using a Bonferroni correction, and are indicated by letters. (**B**) ChIP-seq results of MvaT-V5 and MvaV-V5 over the R-tailocin gene cluster of CHA0 under non-induced (blue) and induced (9 µM mitomycin C, green) conditions. Genes encoding the structural parts of the R-tailocin #1 and the R-tailocin #2 are colored red and green, respectively, the *prtR1* gene is colored pink, the gene encoding the lytic gene holin (*hol*) is colored yellow, other genes belonging to the cluster are colored white, and the bacterial genes neighboring the R-tailocin gene cluster are colored black.

**S12 Fig. MvaT and MvaV may play a role in regulating the expression of the other viral particle clusters in CHA0, while PrtR1 does not bind to these clusters.** Chromatin immunoprecipitation sequencing (ChIP-seq) results of PrtR1-V5, MvaT-V5 and MvaV-V5 over the siphovirus prophage cluster, the myovirus prophage cluster and the cluster in *P. protegens* CHA0 corresponding to the defective prophage 5 in the genome of *Pseudomonas protegens* Pf-5 (2) under non-induced (NI, blue) and induced (I, green, 9 µM mitomycin C) conditions. Genes encoding myovirus or siphovirus structural components are colored purple or yellow, respectively. Genes encoding putative regulatory proteins are colored pink, and genes encoding the t-RNA are colored green.

**S13 Fig. MvaT and MvaV bind to several gene clusters involved in competitive traits in CHA0, including VgrG modules associated with the type VI secretion system (T6SS), CupB fimbriae and biosynthesis of the antibiotic 2,4-diacetylphloroglucinol (DAPG).** Chromatin immunoprecipitation sequencing (ChIP-seq) results of PrtR1-V5, MvaT-V5 and MvaV-V5 over the T6SS VgrG1a module, VgrG1b module, and a putative VgrG module, as well as the biosynthetic clusters for the CupB fimbriae and DAPG under non-induced (blue) and induced conditions (green, 9 µM mitomycin C).

**S14 Fig. MvaT and MvaV bind to some but not all of the gene clusters encoding lipopolysaccharide (LPS) O-antigens in CHA0.** Chromatin immunoprecipitation sequencing (ChIP-seq) results of PrtR1-V5, MvaT-V5 and MvaV-V5 over the OBC1, OBC2, OBC3 (O-PS biosynthesis clusters) and OSA (O-specific antigen) clusters under non-induced (blue) and induced conditions (green, 9 µM mitomycin C).

**S15 Fig. MvaT and MvaV of CHA0 bind to the promoter regions of their own genes, suggesting autoregulatory mechanisms.** Chromatin immunoprecipitation sequencing (ChIP-seq) results of MvaT-V5 and MvaV-V5 over the *mvaT* (**A**) and *mvaV* (**B**) loci under non-induced (NI, blue) and induced (I, green, 9 µM mitomycin C) conditions.

**S16 Fig. Immunoblot showing the presence of the different V5-tagged proteins, prior to induction and after 1 h, 2 h and 3 h of induction with mitomycin C.** To select the time point for collecting the DNA for the ChIP-seq analysis an immunoblot analysis of CHA0 wild type, MvaV-V5, MvaT-V5, PrtR-V5 and ParB-V5 was performed. Samples were collected without induction or 1 h, 2 h and 3 h post induction with 9 µM mitomycin C. Samples were prepared according to the protocol of (3). Samples were run on an acrylamide gel (12 % running gel and 5 % stacking gel) and the PageRuler (Thermo Scientific™) protein ladder was used. Gels were revealed using the SuperSignal West Pico PLUS chemiluminescence substrate kit (Thermo Scientific™) and a Fusion FX.

**S1 Table. Plasmids used in this study.**

**S2 Table. CHA0 derivatives and other bacterial strains used in this study.**

**S3 Table. Oligonucleotides used for the construction of the different mutants and plasmids. S4 Table. RNA-sequencing characteristics.**

**S5 Table. Chromatin immunoprecipitation sequencing (ChIP-seq) characteristics.**

