## Supporting information for "A LexA-like repressor and global H-NS-like regulators enable the fine-tuning of R-tailocin expression in environmental *Pseudomonas*"

#### Supplementary Figures

**S1 Fig.** $H_2O_2$ and mitomycin C induce the expression of the R-tailocin gene cluster and impact the growth of CHA0 wild type and H-NS mutants.
**S2 Fig.** Growth ($OD_{600nm}$) and relative fluorescence (RFU) curves of transcriptional reporters of R-tailocin gene cluster and *prtR1* expression in CHA0 wild type and mutants Δ*prtR1**, Δtailcluster, and Δ*myo*Δ*siph*.
**S3 Fig.** Effect of the deletion of genes with suspected regulatory function on the expression of the R-tailocin gene cluster and *prtR1* in CHA0.
**S4 Fig.** Growth kinetics of CHA0 derivatives expressing the different V5-tagged proteins (MvaT-V5, MavV-V5, PrtR1-V5 and ParB-V5).
**S5 Fig.** ParB binds specifically to regions near the origin and terminus of replication of CHA0.
**S6 Fig.** DNA encoding ribosomal RNA is a highly abundant contaminant in all conditions of the ChIP-seq including the condition without V5-flagged proteins (CHA0 wild type).
**S7 Fig.** The deletion of *mvaT* or *mvaV* leads to the differential regulation of prophage clusters as well as gene clusters related to cell surface decorations and competition traits in CHA0.
**S8 Fig.** The deletion of both *mvaT* and *mvaV* leads to a higher expression of genes related to translation compared to the CHA0 wild type.
**S9 Fig.** Growth ($OD_{600nm}$) and relative fluorescence (RFU) curves of transcriptional reporters of R-tailocin gene cluster and *prtR1* expression in CHA0 wild type, single mutants Δ*mvaT*, Δ*mvaV* and double mutant Δ*mvaT*Δ*mvaV*.
**S10 Fig.** Comparison of the ChIP-seq peaks of MvaT and MvaV under non-induced (A) and induced conditions (B).
**S11 Fig.** MvaT and MvaV of CHA0 are global regulators preferentially binding regions with a high A-T percentage but not the R-tailocin gene cluster of CHA0.
**S12 Fig.** MvaT and MvaV may play a role in regulating the expression of the other viral particle clusters in CHA0, while PrtR1 does not bind to these clusters.
**S13 Fig.** MvaT and MvaV bind to several gene clusters involved in competitive traits in CHA0, including VgrG modules associated with the type VI secretion system (T6SS), CupB fimbriae and biosynthesis of the antibiotic 2,4-diacetylphloroglucinol (DAPG).
**S14 Fig.** MvaT and MvaV bind to some but not all of the gene clusters encoding lipopolysaccharide (LPS) O-antigens in CHA0.
**S15 Fig.** MvaT and MvaV of CHA0 bind to the promoter regions of their own genes, suggesting autoregulatory mechanisms.

**S16 Fig.** Immunoblot showing the presence of the different V5-tagged proteins, prior to induction and after 1 h, 2 h and 3 h of induction with mitomycin C.

#### Supplementary Tables

**S1 Table.** Plasmids used in this study.
**S2 Table.** CHA0 derivatives and other bacterial strains used in this study.
**S3 Table.** Oligonucleotides used for the construction of the different mutants and plasmids.
**S4 Table.** RNA-sequencing characteristics.
**S5 Table.** Chromatin immunoprecipitation sequencing (ChIP-seq) characteristics.

#### References

#### Supplementary Figures

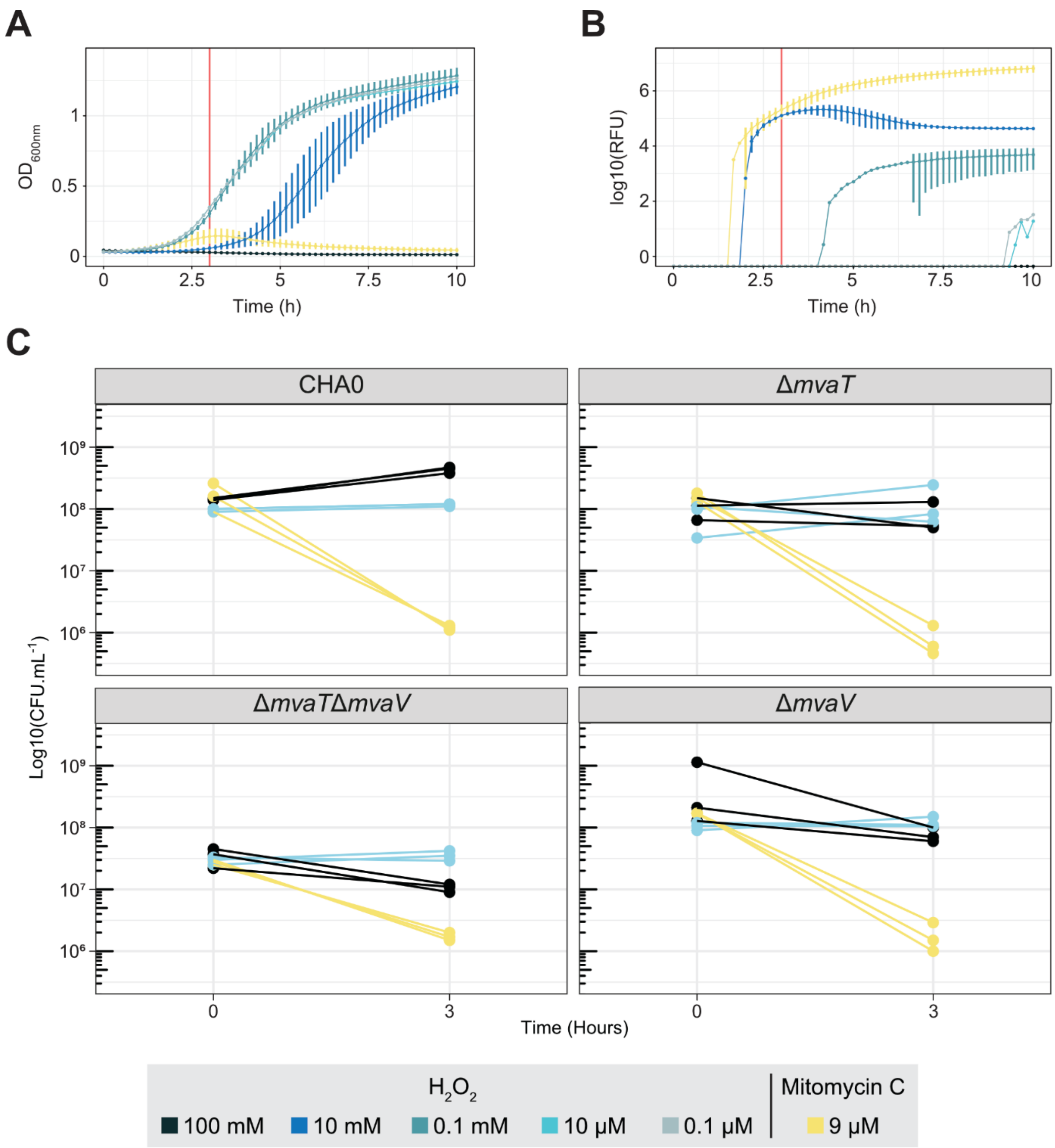


**S1 Fig. $H_2O_2$ and mitomycin C induce the expression of the R-tailocin gene cluster and impact the growth of CHA0 wild type and H-NS mutants.** (**A**, **B**) R-tailocin gene expression was monitored using *P. protegens* CHA0 carrying a transcriptional reporter vector with a copy of the R-tailocin gene cluster promoter located upstream of the *egfp* gene. Optical density at 600 nm (**A**, $OD_{600nm}$) and GFP fluorescence (**B**, relative fluorescence units, RFU) were monitored in rich medium (NYB) following induction with different concentrations of $H_2O_2$ or 9 µM mitomycin C by taking measurements every 10 min in a BioTeK Synergy H1 plate reader. The curves with error bars represent the average (± standard deviation) of measurements from three biological replicates with three technical replicates each. The red bar corresponds to the three-hour time point at which samples were collected for RNA sequencing (RNA-seq). (**C**) Colony forming units (CFU) of the different samples used for the RNA-seq for *P. protegens* CHA0 wild type, Δ*mvaT*, Δ*mvaV* and Δ*mvaT*Δ*mvaV*. Samples were collected prior to induction (at time 0 h) and 3 h after induction with $H_2O_2$ (10 mM), mitomycin C (9 µM) or without induction (control). Three technical replicates were collected for each condition.

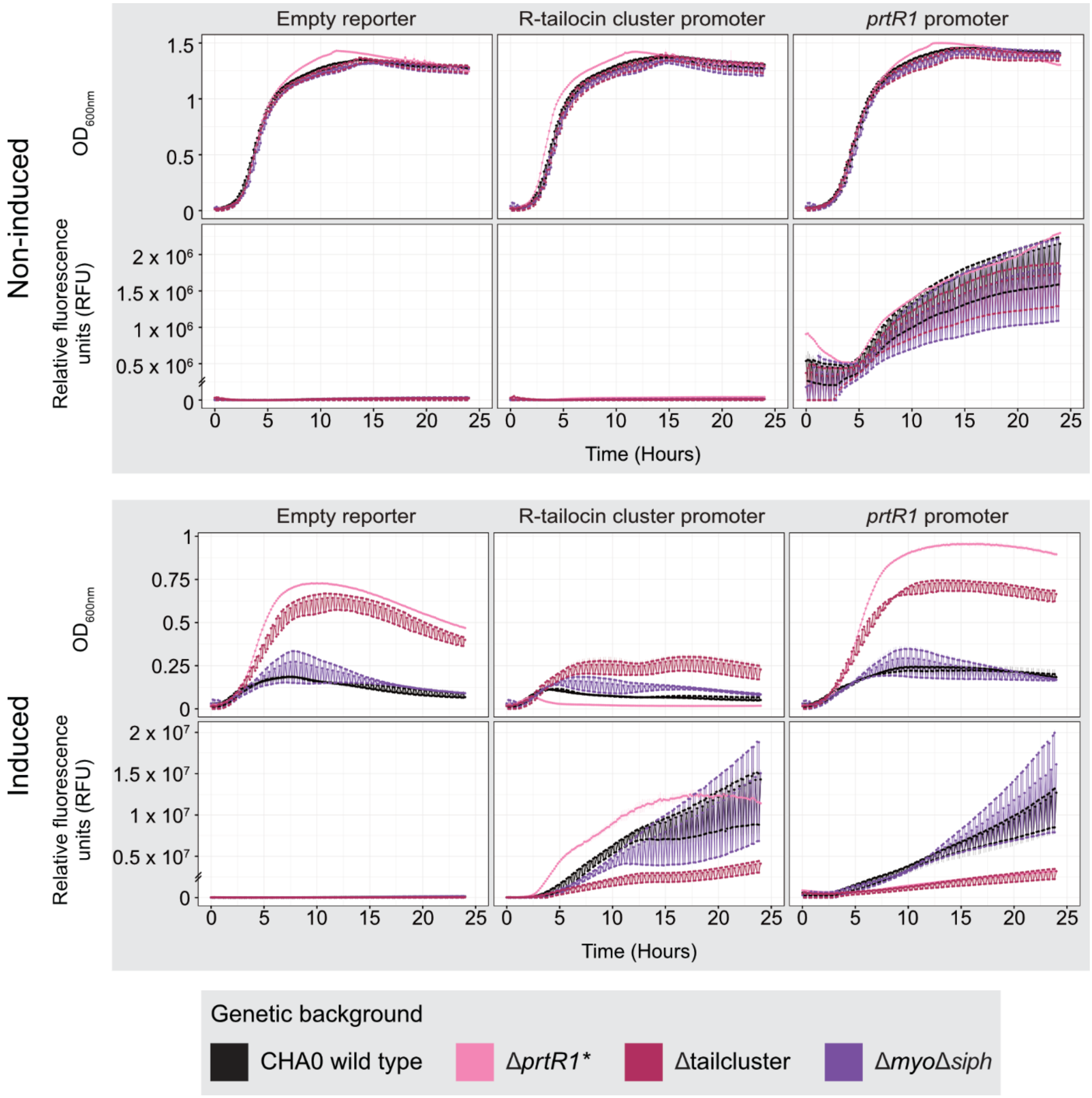


**S2 Fig. Growth ($OD_{600nm}$) and relative fluorescence (RFU) curves of transcriptional reporters of R-tailocin gene cluster and *prtR1* expression in CHA0 wild type and mutants Δ*prtR1**, Δtailcluster, and Δ*myo*Δ*siph*.** The expression of the R-tailocin gene cluster and the locus-specific regulatory gene *prtR1* was monitored using the transcriptional reporters pOT1e-$P_{hol}$-*egfp* and pOT1e-$P_{prtR1}$-*egfp*, respectively, in the wild type CHA0 (black) and its mutants Δ*prtR1** (pink), Δtailcluster (burgundy) and Δ*myo*Δ*siph* (purple). Strains were grown in rich medium (NYB), following induction with 9 µM mitomycin C or without induction. Optical density at 600 nm ($OD_{600nm}$) and GFP fluorescence (relative fluorescence units, RFU) were monitored in rich medium (NYB) every 10 min for 24 h in a BioTeK Synergy H1 plate reader. Curves show means (± standard deviation) of a minimum of three biological replicates with three technical replicates each.

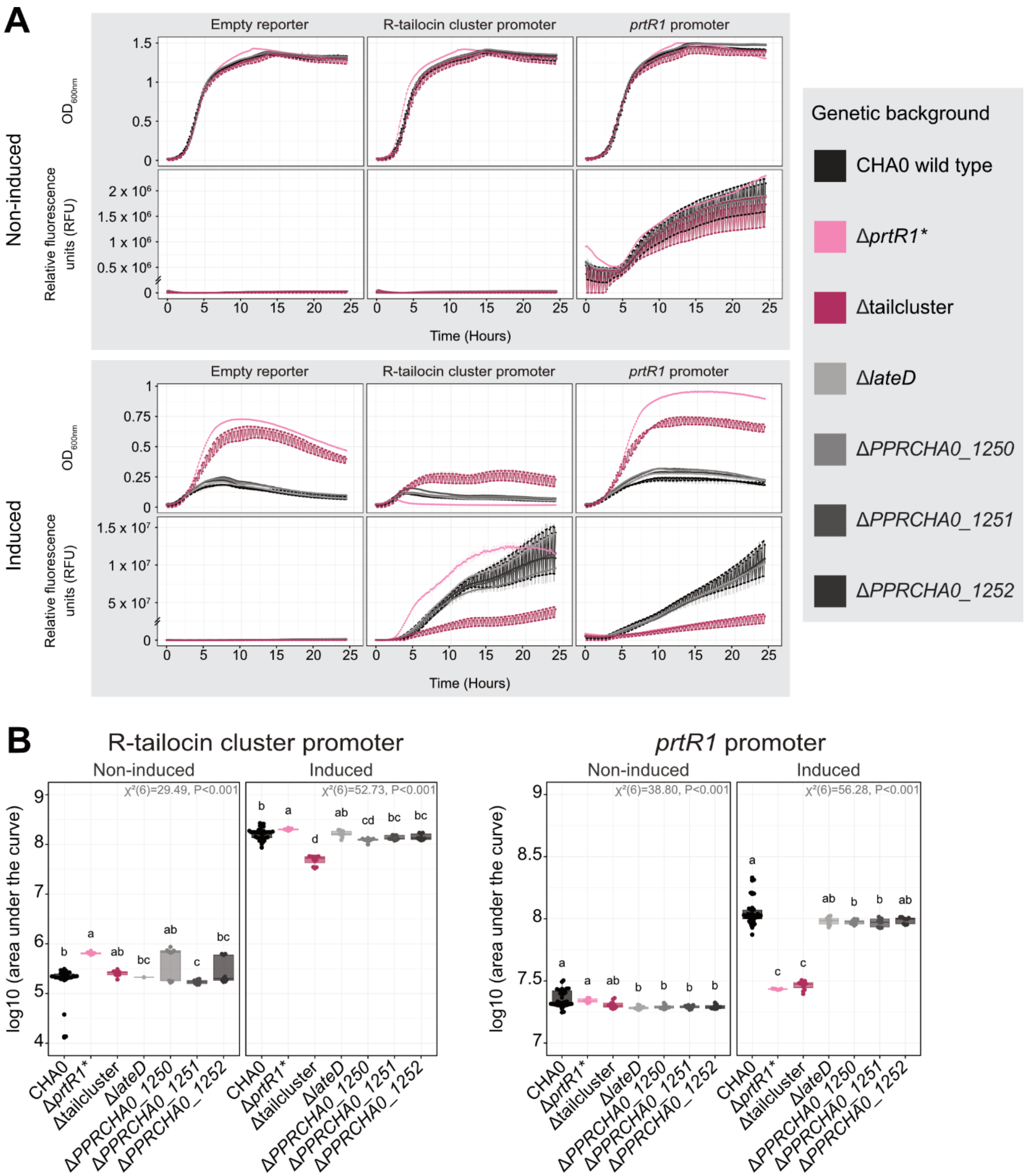


**S3 Fig. Effect of the deletion of genes with suspected regulatory function on the expression of the R-tailocin gene cluster and *prtR1* in CHA0.** The expression was monitored using the transcriptional reporters pOT1e-$P_{hol}$-*egfp* and pOT1e-$P_{prtR1}$-*egfp*, respectively, in the wild type CHA0 (black) and its different mutants (Δ*prtR1**, Δtailcluster, Δ*lateD*, Δ*PPRCHA0_1250*, Δ*PPRCHA0_1251* and Δ*PPRCHA0_1252*). Strains were grown in rich medium (NYB), following induction with 9 µM mitomycin C or without induction. (**A**) Optical density at 600 nm and GFP fluorescence (relative fluorescence units, RFU) were monitored every 10 min for 24 h in a BioTeK Synergy H1 plate reader. (**B**) The data were then used to calculate the area under the RFU curve. Statistical differences were assessed by Kruskal-Wallis tests with Bonferroni correction and are indicated by letters. A minimum of three biological replicates with at least three technical replicates are plotted. Curves show means (± standard deviation) of a minimum of three biological replicates with three technical replicates each.

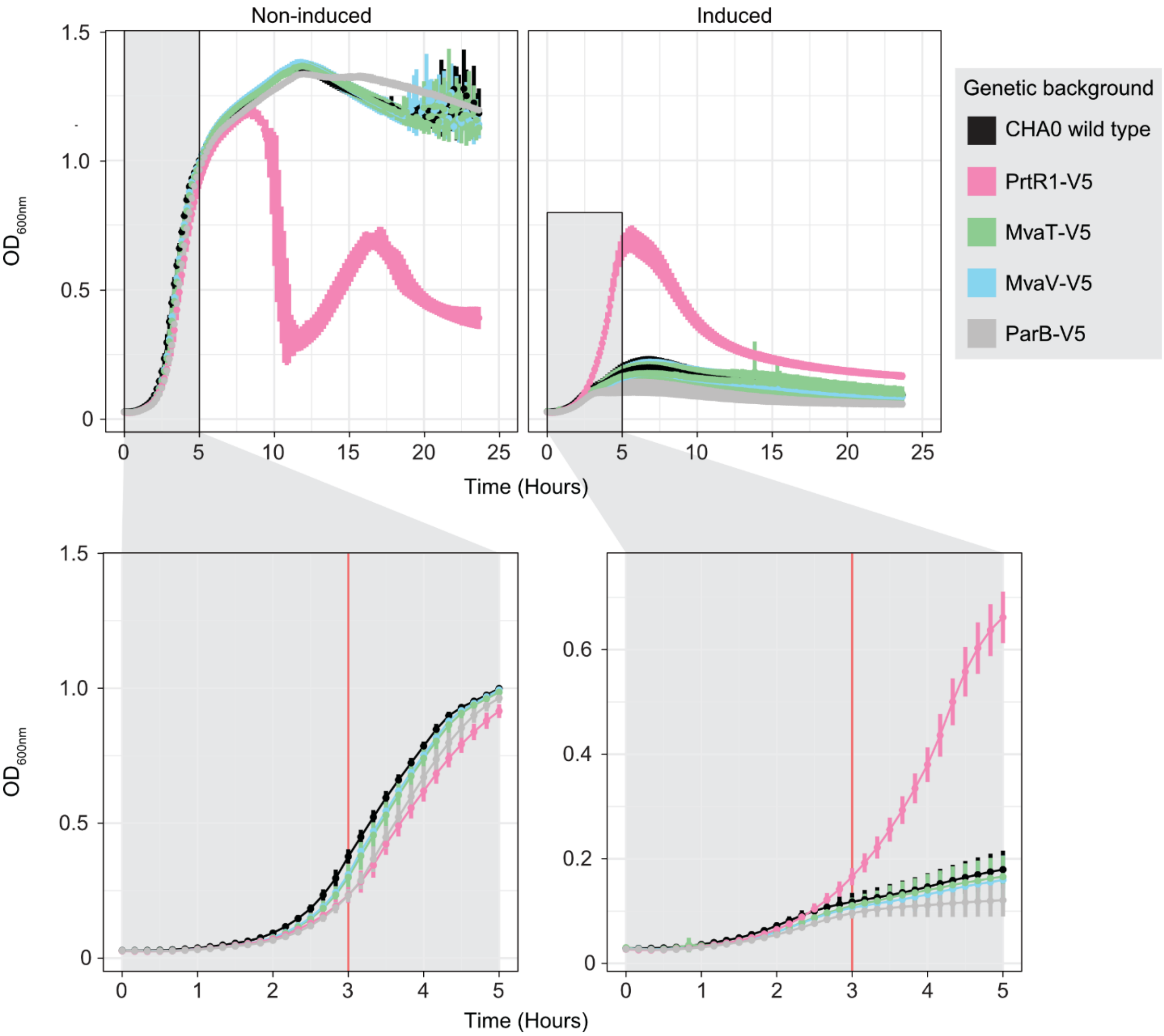


**S4 Fig. Growth kinetics of CHA0 derivatives expressing the different V5-tagged proteins (MvaT-V5, MavV-V5, PrtR1-V5 and ParB-V5).** To test for differences in growth patterns caused by the V5-flagged proteins, the growth of the different strains including the wild type CHA0 in rich medium (NYB) under non-induced and induced (9 µM mitomycin C) conditions was monitored for 24 h by measuring the optical density at 600 nm ($OD_{600nm}$) every 10 min in a BioTeK Synergy H1 plate reader. Curves show means (± standard deviation) of two biological replicates with four technical replicates each. The red bar in the lower panels, representing the first five hours of incubation, corresponds to the three-hour time point at which samples were collected for RNA sequencing.

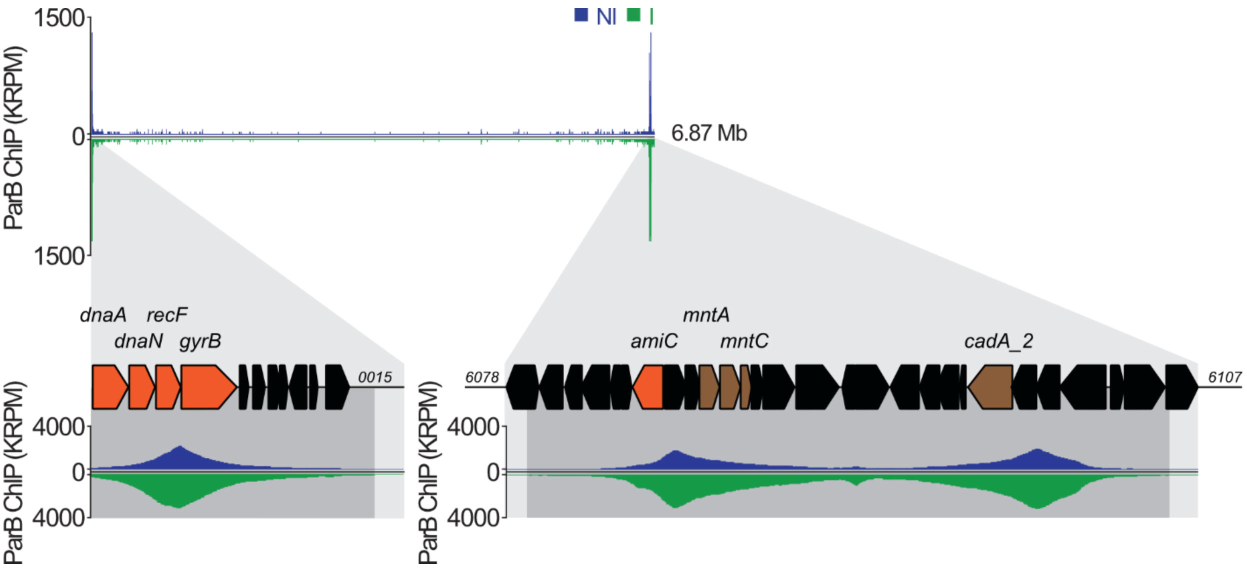


**S5 Fig. ParB binds specifically to regions near the origin and terminus of replication of CHA0.** Chromatin immunoprecipitation sequencing (ChIP-seq) results of ParB-V5 over the origin and terminus of replication under non-induced (NI, blue) and induced (I, 9 µM mitomycin C, green) conditions. The genes involved in replication or DNA repair are colored orange and genes involved in transport are colored brown. Other genes are colored black.

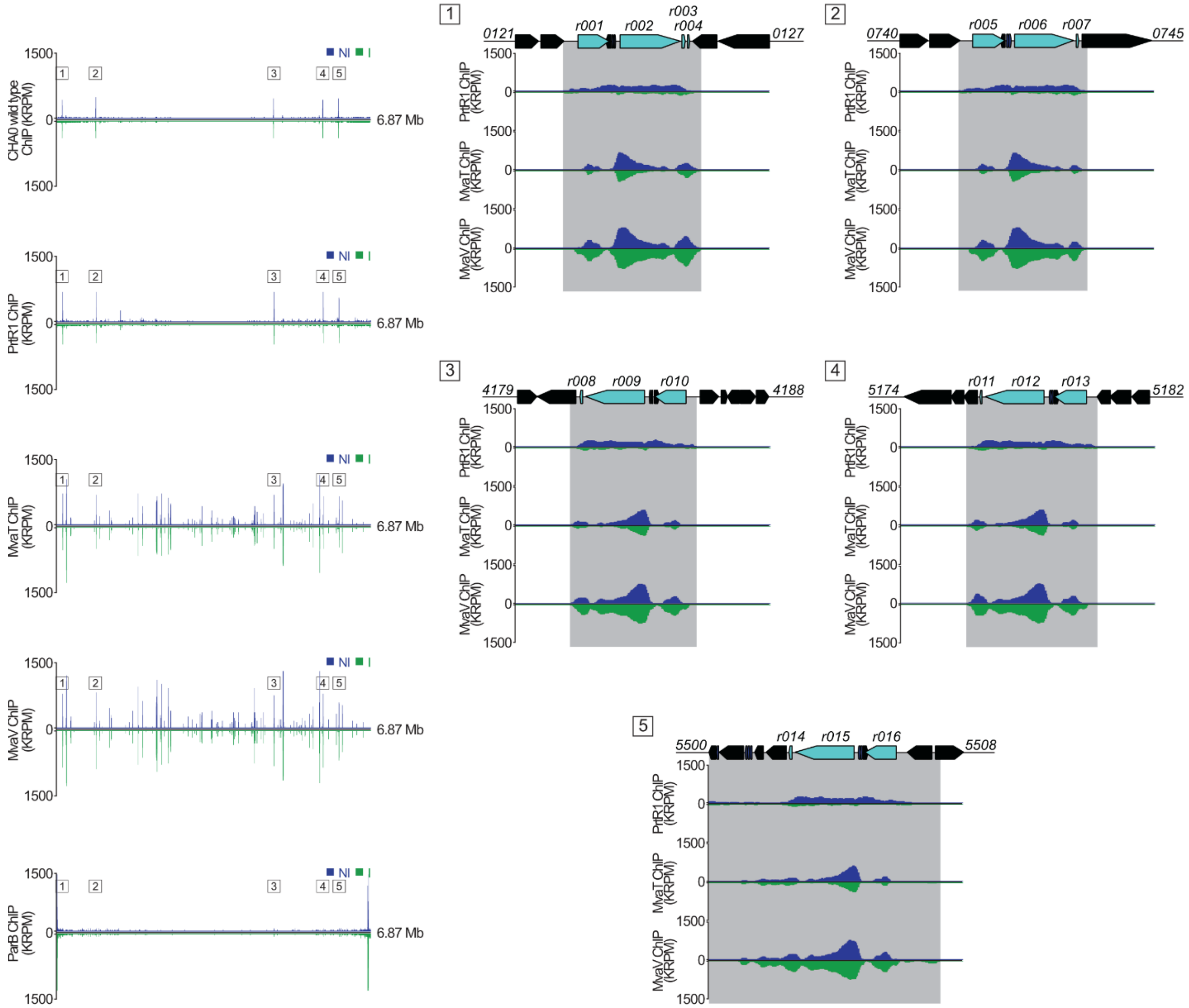


**S6 Fig. DNA encoding ribosomal RNA is a highly abundant contaminant in all conditions of the ChIP-seq including the condition without V5-flagged proteins (CHA0 wild type).** Chromatin immunoprecipitation sequencing (ChIP-seq) for *P. protegens* CHA0 wild type, MvaT-V5, MvaV-V5, PrtR1-V5 and ParB-V5 and with highlighted results of the five different clusters (annotated from 1 to 5) encoding ribosomal RNA in *P. protegens* CHA0, in non-induced (NI, blue) and induced (I, green, 9 µM mitomycin C) conditions. Each cluster encodes at least one 16S, one 23S and one 5S ribosomal subunit. The genes encoding ribosomal RNA and transfer RNA (tRNA) are colored light blue and dark blue, respectively, and other genes are colored black.

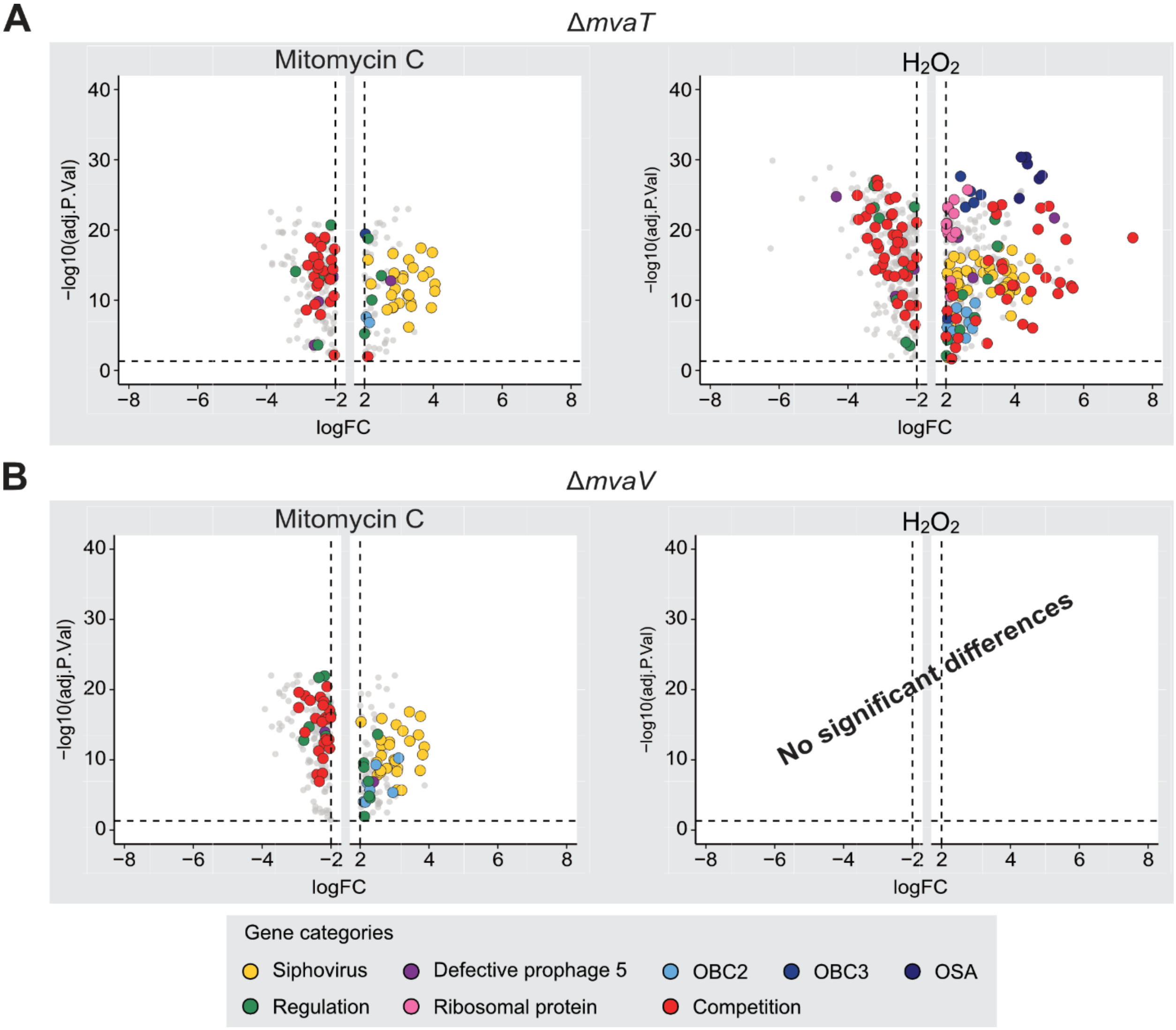


**S7 Fig. The deletion of *mvaT* or *mvaV* leads to the differential regulation of prophage clusters as well as gene clusters related to cell surface decorations and competition traits in CHA0.** Volcano plots showing the RNA sequencing results of the effect of the deletion of *mvaT* (**A**) and *mvaV* (**B**) in *P. protegens* CHA0 following exposure to 9 µM mitomycin C (left) or 10 mM $H_2O_2$ (right). The colored dots correspond to different gene categories, yellow for the siphovirus, purple for the defective prophage 5, blue for the OBC2, OBC3 and OSA clusters, green for regulation, pink for ribosomal proteins and red for competition. The vertical dashed lines correspond to the log2(Fold-change) thresholds set for this analysis (-2 >log2(FC) > 2). The horizontal dashed line corresponds to the significance level of $P < 0.05$.

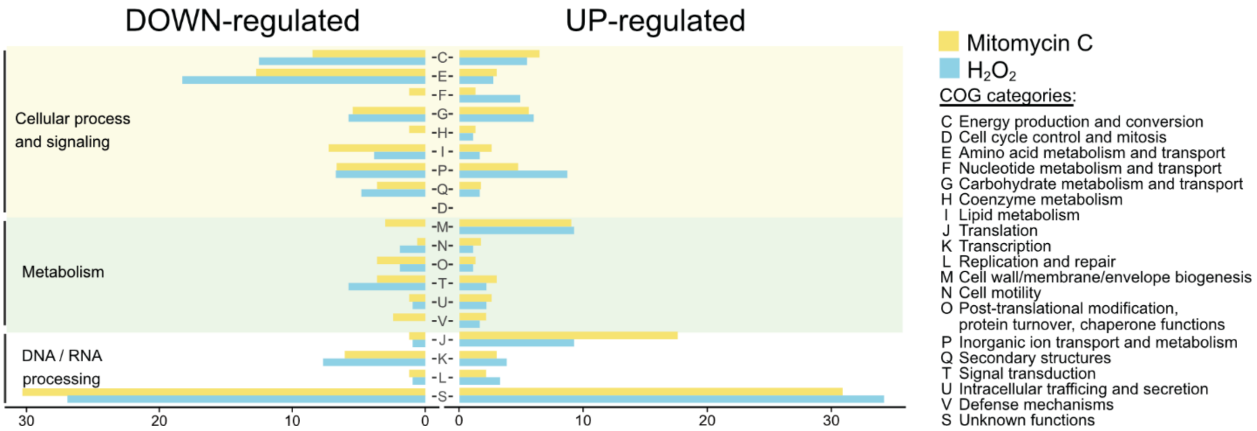


**S8 Fig. The deletion of both *mvaT* and *mvaV* leads to a higher expression of genes related to translation compared to the CHA0 wild type.** Genes that are down-regulated and up-regulated in the Δ*mvaT*Δ*mvaV* mutant compared to the CHA0 wild type according to their COG assignment following exposure to the two compounds. The length of the bars corresponds to the percentage of genes associated with the different COG assignments.

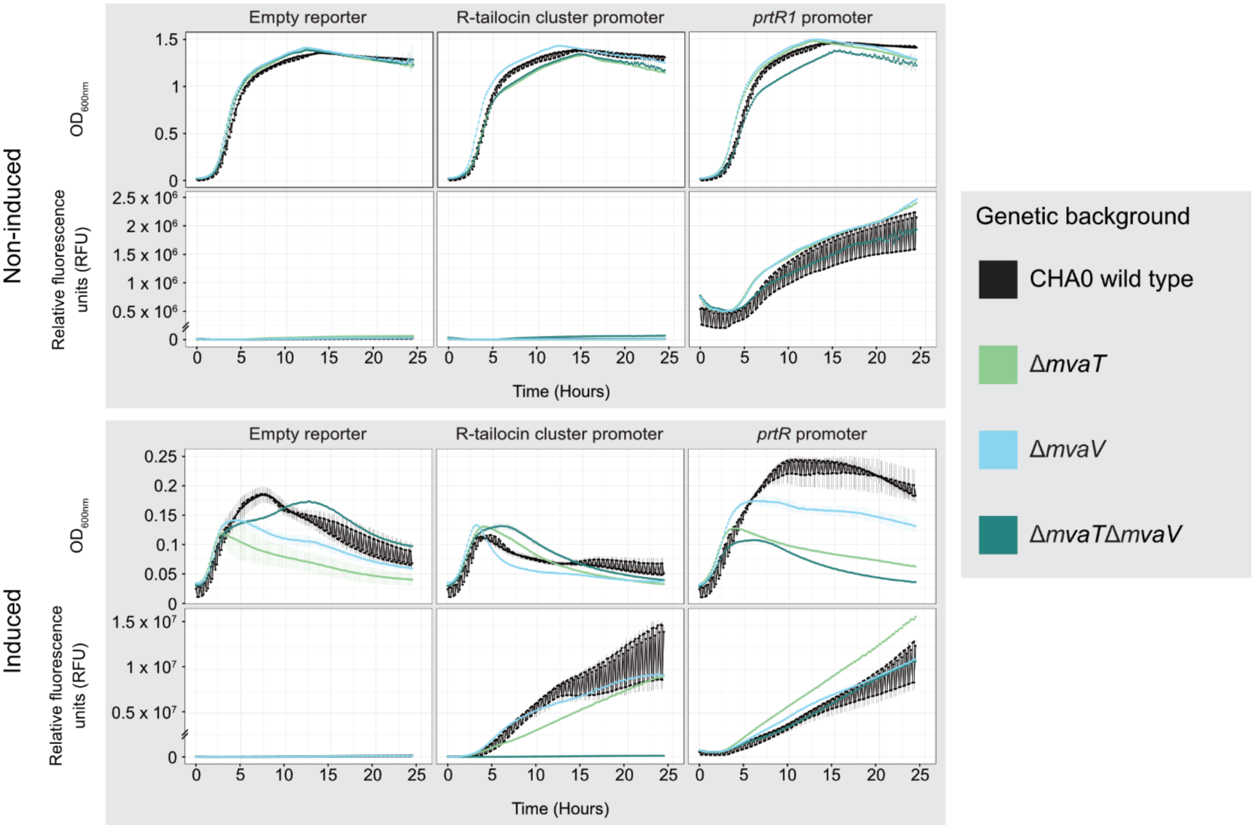


**S9 Fig. Growth ($OD_{600nm}$) and relative fluorescence (RFU) curves of transcriptional reporters of R-tailocin gene cluster and *prtR1* expression in CHA0 wild type, single mutants Δ*mvaT*, Δ*mvaV* and double mutant Δ*mvaT*Δ*mvaV*.** The expression of the R-tailocin gene cluster and the locus-specific regulatory gene *prtR1* was monitored using the transcriptional reporters pOT1e-$P_{hol}$-*egfp* and pOT1e-$P_{prtR1}$-*egfp*, respectively, in the wild type CHA0 (black) and mutants Δ*mvaT*, Δ*mvaV*, Δ*mvaT*Δ*mvaV*. Strains were grown in rich medium (NYB), following induction with 9 µM mitomycin C or without induction. Optical density at 600 nm ($OD_{600nm}$) and GFP fluorescence (relative fluorescence units, RFU) were monitored in rich medium (NYB) every 10 min for 24 h in a BioTeK Synergy H1 plate reader. Curves show means (± standard deviation) of a minimum of three biological replicates with two technical replicates each.

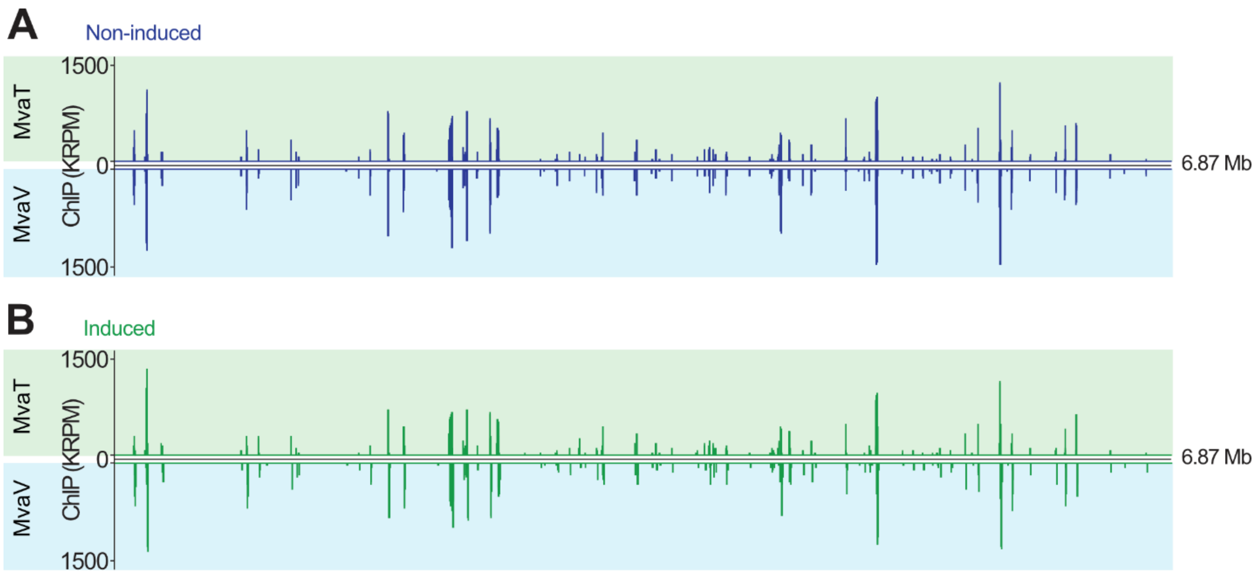


**S10 Fig. Comparison of the ChIP-seq peaks of MvaT and MvaV under non-induced (A) and induced conditions (B).** Chromatin immunoprecipitation sequencing (ChIP-seq) results of MvaT-V5 (green highlight) and MvaV-V5 (blue highlight) under non-induced (blue peaks) and induced (9 µM mitomycin C, green peaks) conditions.

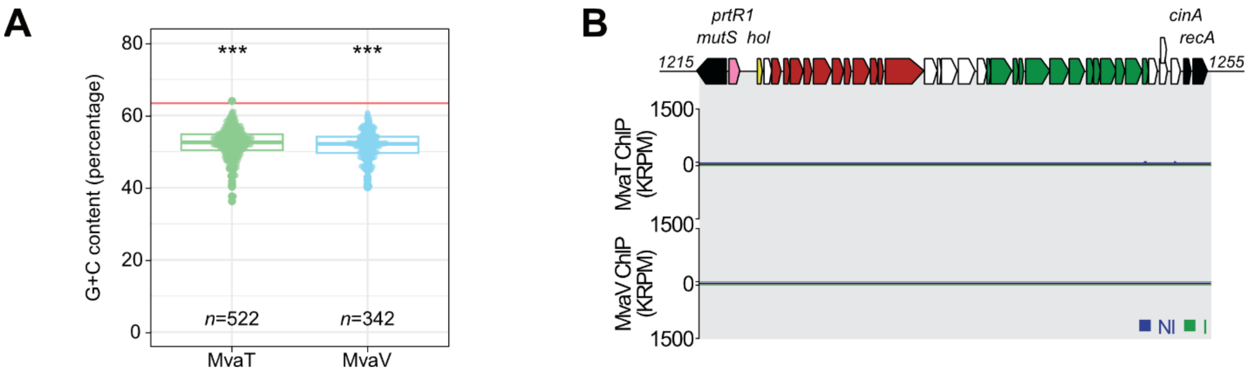


**S11 Fig. MvaT and MvaV of CHA0 are global regulators preferentially binding regions with a high A-T percentage but not the R-tailocin gene cluster of CHA0.** (**A**) GC content of the reads sequenced under the different peaks of MvaT and MvaV. The red vertical line corresponds to the average GC content of the *P. protegens* CHA0 genome (63.4 %). Statistical differences were assessed by Kruskal-Wallis tests coupled with Dunn tests, using a Bonferroni correction, and are indicated by letters. (**B**) ChIP-seq results of MvaT-V5 and MvaV-V5 over the R-tailocin gene cluster of CHA0 under non-induced (blue) and induced (9 µM mitomycin C, green) conditions. Genes encoding the structural parts of the R-tailocin #1 and the R-tailocin #2 are colored red and green, respectively, the *prtR1* gene is colored pink, the gene encoding the lytic gene holin (*hol*) is colored yellow, other genes belonging to the cluster are colored white, and the bacterial genes neighboring the R-tailocin gene cluster are colored black.

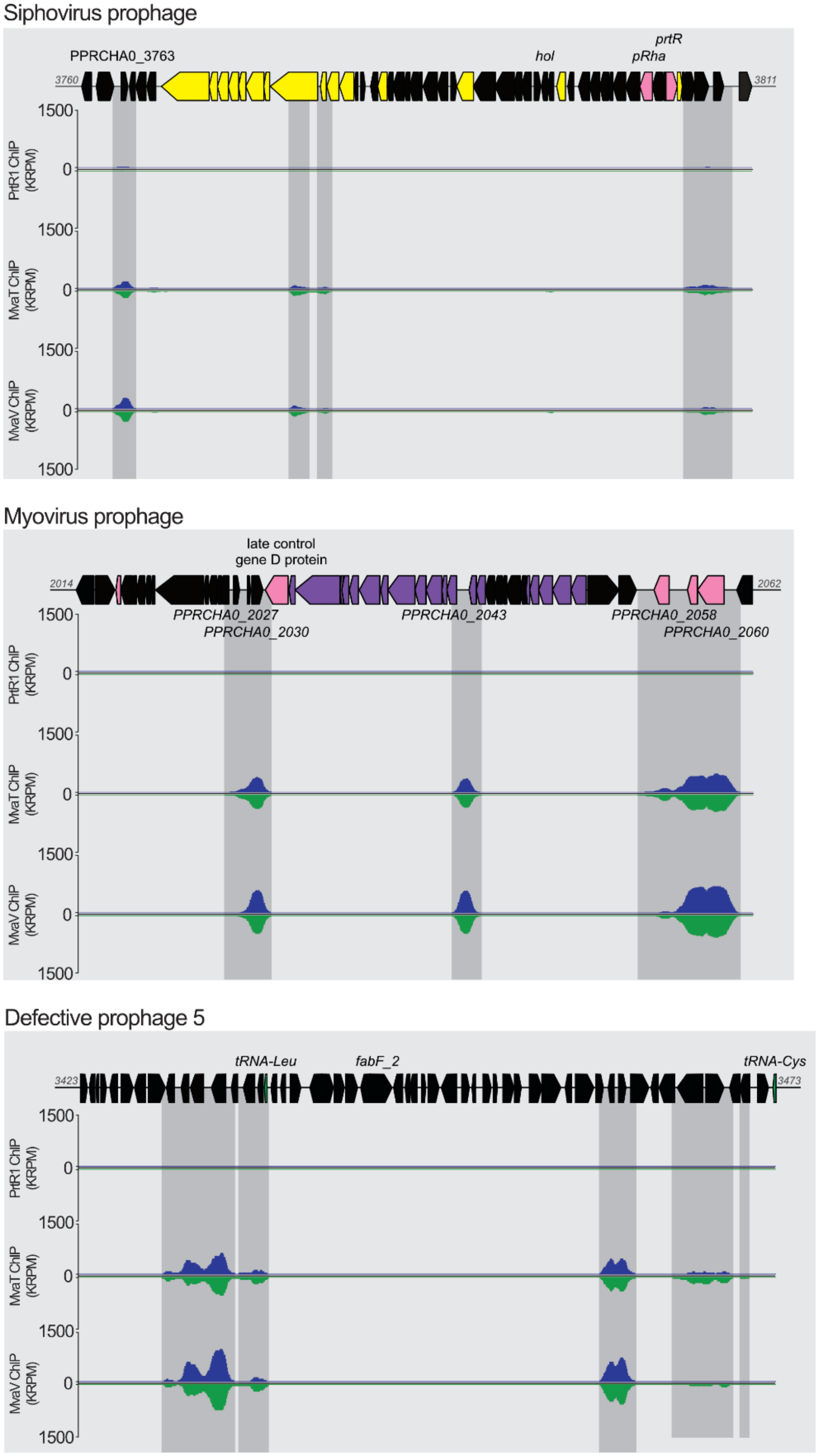


**S12 Fig. MvaT and MvaV may play a role in regulating the expression of the other viral particle clusters in CHA0, while PrtR1 does not bind to these clusters.** Chromatin immunoprecipitation sequencing (ChIP-seq) results of PrtR1-V5, MvaT-V5 and MvaV-V5 over the siphovirus prophage cluster, the myovirus prophage cluster and the cluster in *P. protegens* CHA0 corresponding to the defective prophage 5 in the genome of *Pseudomonas protegens* Pf-5 (2) under non-induced (NI, blue) and induced (I, green, 9 µM mitomycin C) conditions. Genes encoding myovirus or siphovirus structural components are colored purple or yellow, respectively. Genes encoding putative regulatory proteins are colored pink, and genes encoding the t-RNA are colored green.

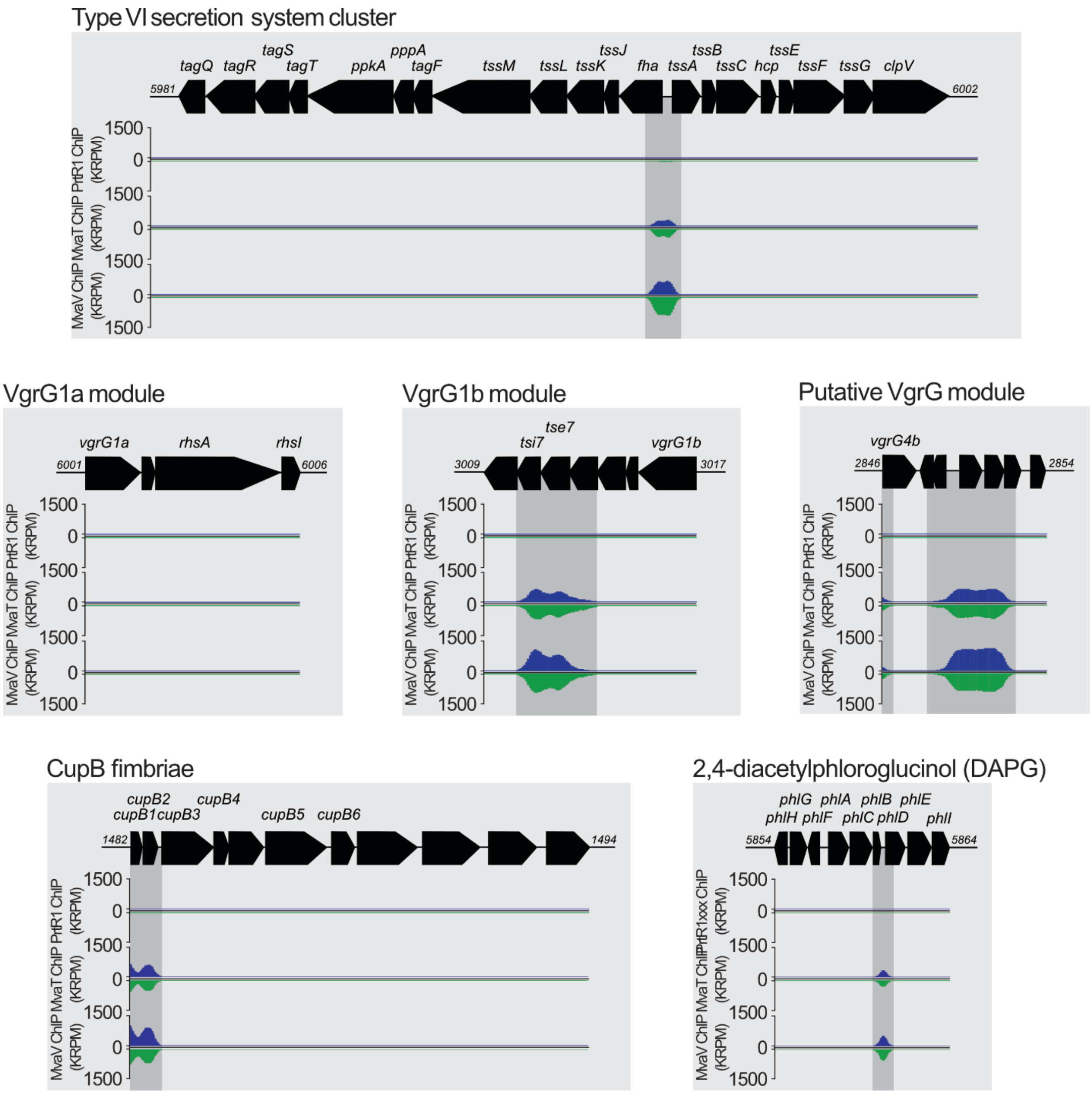


**S13 Fig. MvaT and MvaV bind to several gene clusters involved in competitive traits in CHA0, including VgrG modules associated with the type VI secretion system (T6SS), CupB fimbriae and biosynthesis of the antibiotic 2,4-diacetylphloroglucinol (DAPG).** Chromatin immunoprecipitation sequencing (ChIP-seq) results of PrtR1-V5, MvaT-V5 and MvaV-V5 over the T6SS VgrG1a module, VgrG1b module, and a putative VgrG module, as well as the biosynthetic clusters for the CupB fimbriae and DAPG under non-induced (blue) and induced conditions (green, 9 µM mitomycin C).

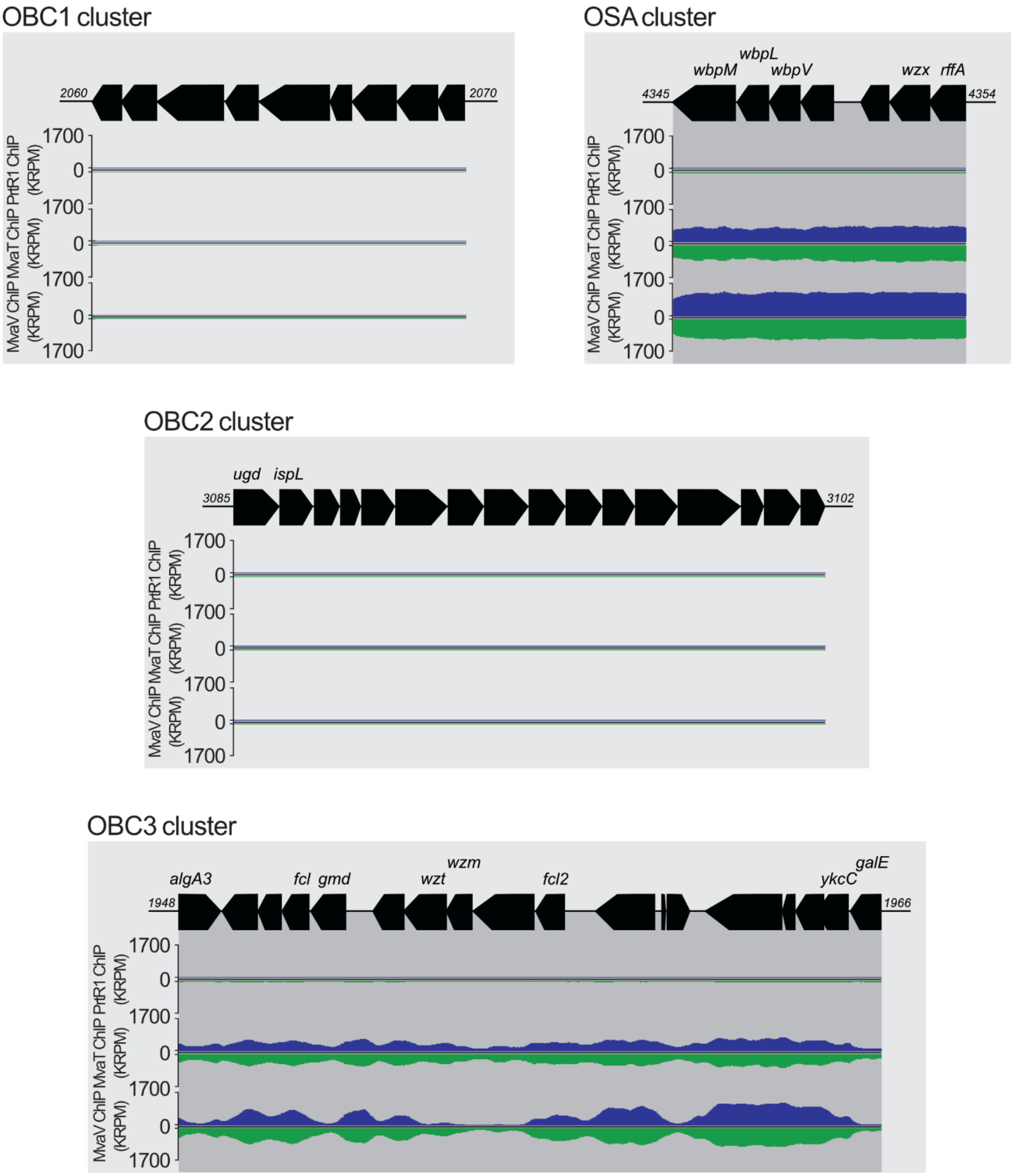


**S14 Fig. MvaT and MvaV bind to some but not all of the gene clusters encoding lipopolysaccharide (LPS) O-antigens in CHA0.** Chromatin immunoprecipitation sequencing (ChIP-seq) results of PrtR1-V5, MvaT-V5 and MvaV-V5 over the OBC1, OBC2, OBC3 (O-PS biosynthesis clusters) and OSA (O-specific antigen) clusters under non-induced (blue) and induced conditions (green, 9 µM mitomycin C).

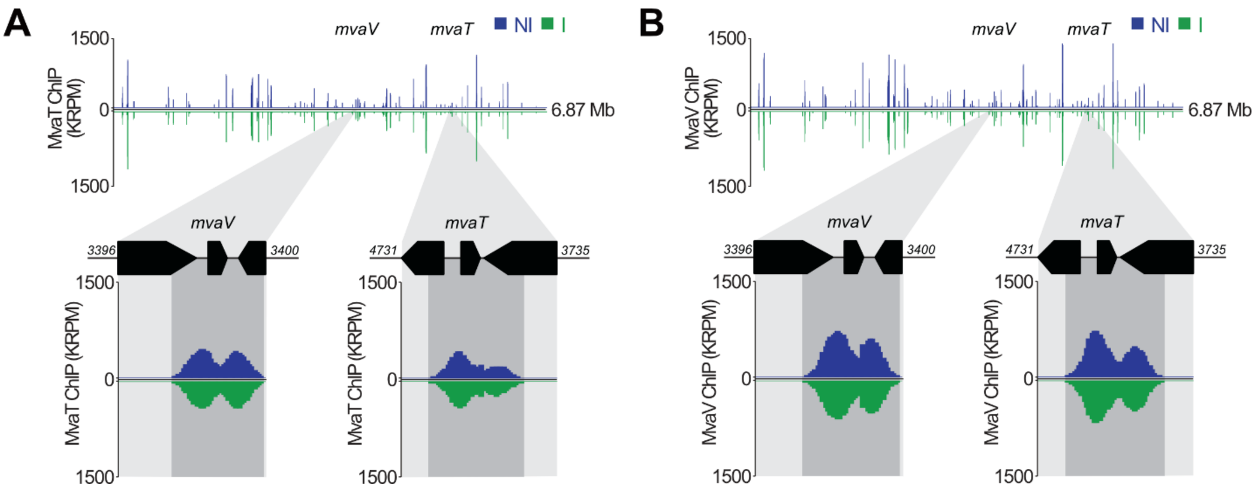


**S15 Fig. MvaT and MvaV of CHA0 bind to the promoter regions of their own genes, suggesting autoregulatory mechanisms.** Chromatin immunoprecipitation sequencing (ChIP-seq) results of MvaT-V5 and MvaV-V5 over the *mvaT* (**A**) and *mvaV* (**B**) loci under non-induced (NI, blue) and induced (I, green, 9 µM mitomycin C) conditions.

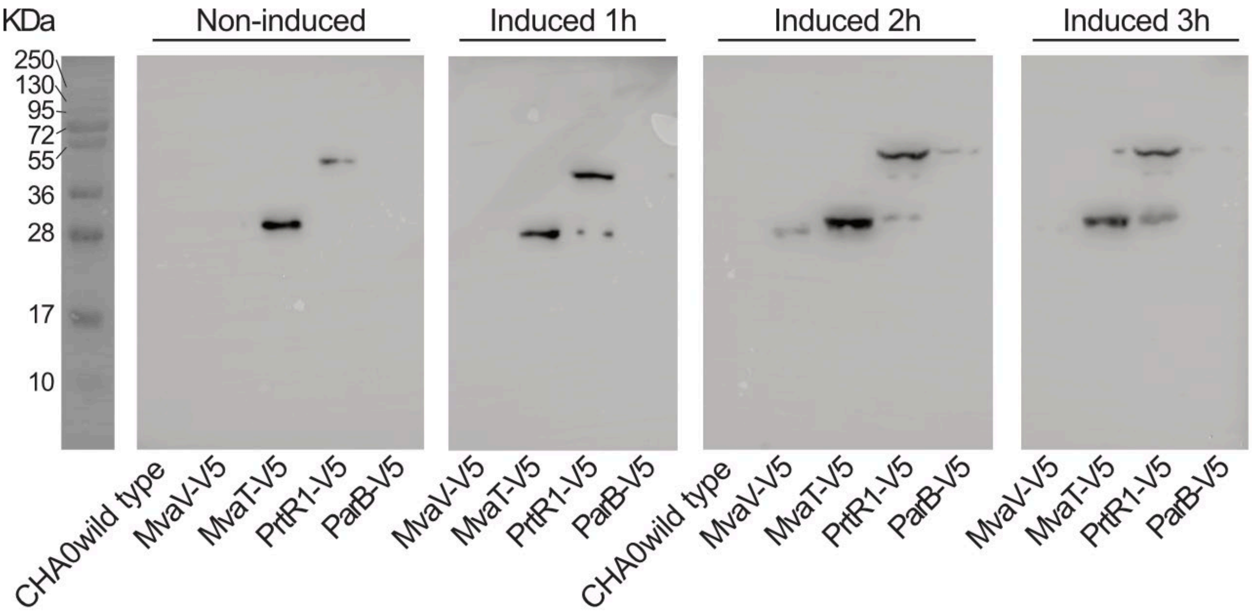


**S16 Fig. Immunoblot showing the presence of the different V5-tagged proteins, prior to induction and after 1 h, 2 h and 3 h of induction with mitomycin C.** To select the time point for collecting the DNA for the ChIP-seq analysis an immunoblot analysis of CHA0 wild type, MvaV-V5, MvaT-V5, PrtR-V5 and ParB-V5 was performed. Samples were collected without induction or 1 h, 2 h and 3 h post induction with 9 µM mitomycin C. Samples were prepared according to the protocol of (3). Samples were run on an acrylamide gel (12 % running gel and 5 % stacking gel) and the PageRuler (Thermo Scientific™) protein ladder was used. Gels were revealed using the SuperSignal West Pico PLUS chemiluminescence substrate kit (Thermo Scientific™) and a Fusion FX.

### Supplementary Tables

##### S1 Table. Plasmids used in this study.

| Plasmid | Genotype or relevant characteristics[1] | Reference or source |
|---|---|---|
| pEMG | Expression vector; *ori*R6K, *lacZα* with two flanking I-SceI sites; $Km^R$, $Ap^R$ | (4) |
| pSW2 | *ori*RK2, *xylS*, *Pm*::*I-SceI*; $Gm^R$ | (4) |
| pME11109 | pEMG::*ΔprtR1*; suicide plasmid for the deletion of the *prtR1* gene; $Km^R$ | This study |
| pME11122 | pEMG::*ΔlateD*; suicide plasmid for the deletion of the *lateD* gene; $Km^R$ | |
| pME11099 | pEMG::*ΔPPRCHA0_1250*; suicide plasmid for the deletion of the PPCHA0_1250 gene; $Km^R$ | This study |
| pME11100 | pEMG::*ΔPPRCHA0_1251*; suicide plasmid for the deletion of the PPCHA0_1251 gene; $Km^R$ | This study |
| pME11095 | pEMG::*ΔPPRCHA0_1252*; suicide plasmid for the deletion of the PPCHA0_1252 gene; $Km^R$ | This study |
| pME11167 | pEMG::mvaT-V5; suicide plasmid for C-terminal tagging of the MvaT protein with V5; $Km^R$ | This study |
| pME11168 | pEMG::mvaV-V5; suicide plasmid for C-terminal tagging of the MvaV protein with V5; $Km^R$ | This study |
| pME11169 | pEMG::prtR1-V5; suicide plasmid for C-terminal tagging of the PrtR1 protein with V5; $Km^R$ | This study |
| pME11170 | pEMG::parB-V5; suicide plasmid for C-terminal tagging of the ParB protein with V5; $Km^R$ | This study |
| pOT1e | Promoter-probe vector based on the pBBR1MCS-5 replicon; contains promoterless *egfp* and MCS between two transcriptional terminators; $Gm^R$ | (5) |
| pOT1e-$P_{hol}$-*egfp** | $P_{hol}$-*egfp* transcriptional fusion in pOT1e; used for monitoring R-tailocin gene cluster expression in CHA0; $Gm^R$ | This study |
| pOT1e-$P_{prtR1}$-*egfp** | $P_{prtR1}$-*egfp* transcriptional fusion in pOT1e; used for monitoring *prtR1* expression in CHA0; $Gm^R$ | This study |

[1] $Ap^R$, ampicillin resistance; $Cm^R$, chloramphenicol resistance; $Gm^R$, gentamicin resistance; $Km^R$, kanamycin resistance.

* See **Figure 2A** for more information on these constructions.

**S2 Table. CHA0 derivatives and other bacterial strains used in this study.**

| Strain name | Strain code | Genotype or relevant characteristics[1] | Reference or source |
|---|---|---|---|
| ***Pseudomonas protegens* CHA0 and derivatives** | | | |
| CHA0 | $CHA0^T$ | *P. protegens* type strain; wild type; genome accession no. LS999205.1 | (6,7) |
| Δ*mvaT* | CHA1121 | Deletion of *mvaT* in CHA0 | (8) |
| Δ*mvaV* | CHA1126 | Deletion of *mvaV* in CHA0 | (8) |
| Δ*mvaT*Δ*mvaV* | CHA1127 | Deletion of *mvaT* and *mvaV* in CHA0 | (8) |
| Δ*prtR1* | - | Attempted deletion of *prtR1* of CHA0; lethal due to de-repression of the entire R-tailocin gene cluster including its cell lysis genes | This study |
| Δtailcluster | CHA5285 | Deletion of the entire R-tailocin gene cluster of CHA0 [from PPRCHA0_1217 to PPRCHA0_1252] | (9) |
| Δ*prtR1** | CHA5451 | Deletion of *prtR1* in the CHA0 Δtailcluster mutant | This study |
| Δ*lateD* | CHA5486 | Deletion of *lateD* (PPRCHA0_1234) of the tailocin gene cluster of CHA0 | This study |
| Δ*PPRCHA0_1250* | | Deletion of PPRCHA0_1250 of the tailocin gene cluster of CHA0 | This study |
| Δ*PPRCHA0_1251* | | Deletion of PPRCHA0_1251 of the tailocin gene cluster of CHA0 | This study |
| Δ*PPRCHA0_1252* | | Deletion of PPRCHA0_1252 of the tailocin gene cluster of CHA0 | This study |
| Δ*myo*Δ*siph* | CHA5299 | CHA0 with the deletion of the *Myoviridae* and *Siphoviridae* prophages | (9) |
| *mvaT*-V5 | CHA5469 | CHA0 with a V5 C-terminally-tagged MvaT | This study |
| *mvaV*-V5 | CHA5470 | CHA0 with a V5 C-terminally-tagged MvaV | This study |
| *prtR1*-V5 | CHA5471 | CHA0 with a V5 C-terminally-tagged PrtR1 | This study |
| *parB*-V5 | CHA5472 | CHA0 with a V5 C-terminally-tagged ParB | This study |
| **Other bacterial strains** | | | |
| *Escherichia coli* S17-1/λpir | | Laboratory strain | (10) |
| *Escherichia coli* DH5α | | Laboratory strain | (11) |

[1] $Gm^R$, gentamicin resistance.

**S3 Table. Oligonucleotides used for the construction of the different mutants and plasmids.**

| Primer | Sequence (5'-3') [1] | Application |
|---|---|---|
| prtR1-del-1 | CGGAATTCCGACTCGCCGAGCTTTAC | Deletion of *prtR1* |
| prtR1-del-2 | CGGGATCCCATAGAACGCATATTGCTCGA | Deletion of *prtR1* |
| prtR1-del-3 | CGGGATCCTAAGCCAGCCCTCTTGAA | Deletion of *prtR1* |
| prtR1-del-4 | ACGCGTCGACAAGGGTCCCTGTCTGTTG | Deletion of *prtR1* |
| check-PrtR1-F | GTCGCCCATGCGGTAGAA | Check of Δ*prtR1*Δtailcluster |
| check-PrtR1-R | GGGGGCGCATAAACAAAG | Check of Δ*prtR1*Δtailcluster |
| del-PPRCHA0_1250-1 | CGGAATTCTATCACGCATCGTTCGTG | Deletion of PPRCHA0_1250 |
| del-PPRCHA0_1250-2 | GGGGTACCCATGGTCAGTGTTTGCAGGTC | Deletion of PPRCHA0_1250 |
| del-PPRCHA0_1250-3 | GGGGTACCTGTTTGCCGGTGGTGTGA | Deletion of PPRCHA0_1250 |
| del-PPRCHA0_1250-4 | CGGGATCCCAGCAACATCTCCGTAGT | Deletion of PPRCHA0_1250 |
| check-PPRCHA0_1250-F | GTTACTGCAACTTGCGGT | Check of ΔPPRCHA0_1250 |
| check-PPRCHA0_1250-R | TCACTCCTCCCTTCCACC | Check of ΔPPRCHA0_1250 |
| del-PPRCHA0_1251-1 | CGGAATTCCTTGCTCAGATCGGACAC | Deletion of PPRCHA0_1251 |
| del-PPRCHA0_1251-2 | GGGGTACCCATCAAGCCCTCCTGAGG | Deletion of PPRCHA0_1251 |
| del-PPRCHA0_1251-3 | GGGGTACCCCCTGAGAGCAGGTTGTC | Deletion of PPRCHA0_1251 |
| del-PPRCHA0_1251-4 | ACGCGTCGACGCCCGGAGCGATAACAAA | Deletion of PPRCHA0_1251 |
| check-PPRCHA0_1251-F | CAGATCGATACGCCGAAG | Check of ΔPPRCHA0_1251 |
| check-PPRCHA0_1251-R | TACCTCACCAACGACCGG | Check of ΔPPRCHA0_1251 |
| del-PPRCHA0_1252-1 | CGGAATTCGATAGAGCCTGCGTTGTG | Deletion of PPRCHA0_1252 |
| del-PPRCHA0_1252-2 | GGGGTACCCATTCCGAGCGTACATGC | Deletion of PPRCHA0_1252 |
| del-PPRCHA0_1252-3 | GGGGTACCTGACGTGCCTCTATCAGT | Deletion of PPRCHA0_1252 |
| del-PPRCHA0_1252-4 | ACGCGTCGACGACCACTTCACGACTGAC | Deletion of PPRCHA0_1252 |
| check-PPRCHA0_1252-F | AAGGTCTATCAGACGGGC | Check of ΔPPRCHA0_1252 |
| check-PPRCHA0_1252-R | AAAATGCGCATGACTCCT | Check of ΔPPRCHA0_1252 |
| check mvaT-V5_F | ATGGGCGAGTACTCCAAA | Check of *mvaT*-V5 |
| check mvaT-V5_R | CTGGATCAATGGCAGGAC | Check of *mvaT*-V5 |
| check mvaV-V5_F | CGTGACATCATCGCCATC | Check of *mvaV*-V5 |
| check mvaV-V5_R | TCTGCGGGTTACCCATTC | Check of *mvaV*-V5 |
| check prtR1-V5_F | CAGCTGTATCGCCTACCT | Check of *prtR1*-V5 |
| check prtR1-V5_R | ATTCATCGCTGCCTTTGC | Check of *prtR1*-V5 |
| check parB-V5_F | CATCGCTCTTGCTCCTCA | Check of *parB*-V5 |
| check parB-V5_B | CTCGGATTGCCGGAAAAT | Check of *parB*-V5 |
| exp_hol-F | GGGGTACCGCCAGCCCTCTTGAACAAAG | Amplification of the promoter region of *hol* to construct pOT1e-$P_{hol}$-*egfp* |
| exp_hol-R | CGGAATTCTCACGTCTCCCGTGCGGATT | Amplification of the promoter region of *hol* to construct pOT1e-$P_{hol}$-*egfp* |
| exp_prtR_F | CCCAAGCTTAGGCGCTACTGTGTTTGT | Amplification of 50 bp in the promoter region of *prtR1* to construct pOT1e-$P_{prtR1}$-*egfp* |
| exp_prtR_R | CACTAGTGCTGTTTTGCATATTTGTGCT | Amplification of 50 bp in the promoter region of *prtR1* to construct pOT1e-$P_{prtR1}$-*egfp* |
| M13_F | GTAAAACGACGGCCAGT | Sequencing verification of the pEMG-based plasmids |
| M13_R | AACAGCTATGACCATG | Sequencing verification of the pEMG-based plasmids |
| pOT1e_F | CCGGTGGATGACCTTTTG | Sequencing verification of the pOT1e -based plasmids |
| pOT1e_R | ACCCTCTCCACTGACAGA | Sequencing verification of the pOT1e -based plasmids |

[1] Restriction sites are underlined.

**S4 Table. RNA-sequencing characteristics.**

| Sample | Replicate | Total number of reads obtained | Number of cleaned reads | Total of HQ mapped reads | Percentage of HQ mapped reads | Mean read length (bp) |
|---|---|---|---|---|---|---|
| Control | 1 | 23,884,484 | 13,778,603 | 13,696,068 | 99.400992 | 135.61795 |
| Control | 2 | 28,770,435 | 20,126,476 | 19,897,262 | 98.861132 | 142.69819 |
| Control | 3 | 30,289,575 | 19,965,886 | 19,739,544 | 98.866356 | 143.0677 |
| Control | 4 | 28,945,914 | 18,533,435 | 18,334,933 | 98.928952 | 142.25514 |
| CHA0 x $H_2O_2$ | 1 | 28,719,152 | 18,573,805 | 18,408,724 | 99.111216 | 143.57343 |
| CHA0 x $H_2O_2$ | 2 | 28,563,233 | 18,970,239 | 18,699,265 | 98.571584 | 143.91466 |
| CHA0 x $H_2O_2$ | 3 | 31,701,482 | 22,255,234 | 22,090,516 | 99.259868 | 142.11202 |
| CHA0 x $H_2O_2$ | 4 | 34,934,772 | 25,469,158 | 25,247,099 | 99.128126 | 144.52022 |
| CHA0 x MMC | 1 | 27,800,039 | 16,839,143 | 16,617,562 | 98.684131 | 143.73303 |
| CHA0 x MMC | 2 | 30,815,428 | 18,836,469 | 18,390,562 | 97.632746 | 142.31945 |
| CHA0 x MMC | 3 | 28,184,857 | 18,198,636 | 18,073,283 | 99.311196 | 142.77518 |
| CHA0 x MMC | 4 | 31,160,531 | 20,328,441 | 20,159,819 | 99.170512 | 141.29277 |

**S5 Table. Chromatin immunoprecipitation sequencing (ChIP-seq) characteristics.**

| Sample | Induction | Sequencing depth | Mapped fragment number | Alignment rate | Duplication rate | Estimated Library size |
|---|---|---|---|---|---|---|
| CHA0 | NI | 39,749,764 | 32,442,223 | 81.62 % | 94.72 % | 1,713,664 |
| CHA0 | I | 39,749,764 | 41,352,762 | 82.48 % | 92.98 % | 2,902,024 |
| MvaT-V5 | NI | 50,829,796 | 47,896,460 | 94.23 % | 52.69 % | 28,207,819 |
| MvaT-V5 | I | 47,294,867 | 45,536,697 | 96.28 % | 60.62 % | 20,293,066 |
| MvaV-V5 | NI | 71,868,578 | 66,791,702 | 92.94 % | 53.92 % | 37,630,616 |
| MvaV-V5 | I | 50,844,011 | 47,988,081 | 94.38 % | 50.56 % | 30,434,888 |
| PrtR1-V5 | NI | 52,027,918 | 50,427,699 | 96.92 % | 66.23 % | 18,351,590 |
| PrtR1-V5 | I | 45,353,799 | 43,904,909 | 96.81 % | 58.24 % | 21,278,060 |
| ParB-V5 | NI | 51,692,357 | 49,460,393 | 95.68 % | 76.26 % | 11,971,155 |
| ParB-V5 | I | 42,109,981 | 40,432,691 | 96.02 % | 87.35 % | 5,119,295 |
